# CRD: a *De novo* Design algorithm for prediction of Cognate Protein Receptors for small molecule ligands

**DOI:** 10.1101/2023.03.30.534983

**Authors:** Santhosh Sankar, Nagasuma Chandra

## Abstract

While predicting a new ligand to bind to a protein is possible with current methods, the converse of predicting a receptor for a ligand is highly challenging, except for very closely-related known protein-ligand complexes. Predicting a receptor for any given ligand will be path-breaking in understanding protein function, mapping sequence-structure-function relationships and for several aspects of drug discovery including studying the mechanism of action of phenotypically discovered drugs, off-target effects and drug repurposing. We use a novel approach for predicting receptors for a given ligand through *de novo* design combined with structural bioinformatics. We have developed a new algorithm CRD, that has multiple modules which combines fragment-based sub-site finding, a machine learning function to estimate the size of the site, a genetic algorithm that encodes knowledge on protein structures and a physics-based fitness scoring scheme. CRD has a pseudo-receptor design component followed by a mapping component to identify possible proteins that house the site. CRD is designed to cater to ligands with known and unknown complexes. CRD accurately recovers sites and receptors for several known natural ligands including ATP, SAM, Glucose and FAD. It designs similar sites for similar ligands, yet to some extent distinguishes between closely related ligands. More importantly CRD correctly predicts receptor classes for several drugs such as penicillins and NSAIDs. We expect CRD to be a valuable tool in fundamental biology research as well as in the drug discovery and biotechnology industry.

## Introduction

Protein function is critically dependent on ligand recognition, and there is a wealth of data on biochemical aspects as well as three-dimensional structures of protein-ligand complexes [1–3], which provides a fundamental critical layer of information for drug discovery and a range of other applications. While there are many tools to aid in the prediction of a ligand that can bind to a given protein [4–6], the converse, that is to predict a protein that can bind to a given ligand remains a daunting problem. If a protein that a ligand can bind to can be identified, it would greatly improve our understanding on how proteins recognize ligands and take us a step closer towards a first-principle understanding of the effect of small molecules on biological systems. More importantly, it will bridge a large gap in drug discovery pipelines, by predicting receptors for candidate drug-libraries, phenotypically discovered drugs thereby facilitating in unraveling the mechanism of drug action and identifying off-target effects.

Traditional approaches for identifying receptors are based on the premise that similar proteins recognize similar ligands and use similarities in the ligand structures in known protein-bound complexes to extrapolate to an unknown ligand [7]. They are severely limited by requiring prior knowledge of protein-ligand complexes whose ligands share significant similarity with the query ligand. A rational de novo approach promises to overcome this limitation and provide a general framework for identifying a protein to bind any given ligand. De novo methods have the distinct advantage of ‘reasoning from first principles’, and provide solutions even when prior examples or reference cases are not available [8,9]. In recent years, de novo approaches have been successfully applied in several areas in biology and biomedical sciences, such as de novo assembly of genome sequences and de novo drug design [10–12]. In the field of proteins, notable examples are the Rosetta suite of tools for protein design with tailored functionalities [13], modifying natural binding sites to improve affinity of known ligands or to recognize different ligands [14,15]. Most of these methods require protein scaffolds in order to place, mutate, and optimize amino acids [14–16]. Currently, there is no method for predicting proteins for ligands that have no reference cases of protein complexes.

In this work, we employ a ‘design’ strategy and develop a de novo algorithm (CRD) for cognate receptor discovery. The term receptor is defined broadly as a protein that recognizes the ligand specifically to perform its function. CRD designs sub-structures mimicking binding sites for a given ligand and identifies proteins that house such binding sites. De novo design of binding sites is a multi-step complex process faced with many challenges and is computationally challenging. In developing CRD, we address the key problems at each step starting from ligand fragmentation to evaluation of generated sites, which we believe will be useful as stand-alone modules for a variety of applications.

## Results

### CRD design strategy

CRD is a novel design-based approach for predicting a receptor for a given ligand. It designs a binding site(s) that can theoretically bind a given ligand, which is scanned against the entire database of known protein-ligand complexes to find a matching site and the receptor protein. It utilizes a fragment-based method involving a combination of chemoinformatics, structural bioinformatics, a genetic algorithm and machine learning to design the sites (Fig. 1). The basic idea is that a given ligand can be deconstructed to a set of fragments that occur in multiple other ligands, whose interaction environment can be learnt from known protein-ligand complexes and applied to designing a binding site for the whole ligand [16]. A main challenge in selection is that the residues need to be selected without any association with a protein structural fold. The algorithm has 3 modules (Fig. 1), each solving several challenges in themselves, in order to get the final outcome. CRD starts with 3D coordinates of a query ligand as an input. The ***FragSite*** module generates fragments of the ligand, compares the fragments to a library of fragments from PDB ligands, aligns the fragments, identifies sites for fragments and generates a residue pool placed on a common 3D framework of the query ligand. The ***SiteDesign*** module generates candidate site residues from large residue-libraries, identifies the size range for the site through probability-based deep learning algorithm, generates seed sites from which *SiteGen*, a Genetic Algorithm selects fittest sites for a given ligand with the help of a triplet-fitness scoring scheme. The ***SiteMapper*** module analyzes the final fittest sites, clusters, identifies number of site types and finds matches if present from PDB sites, thereby identifying possible receptors for a given ligand.

**Fig. 1.**
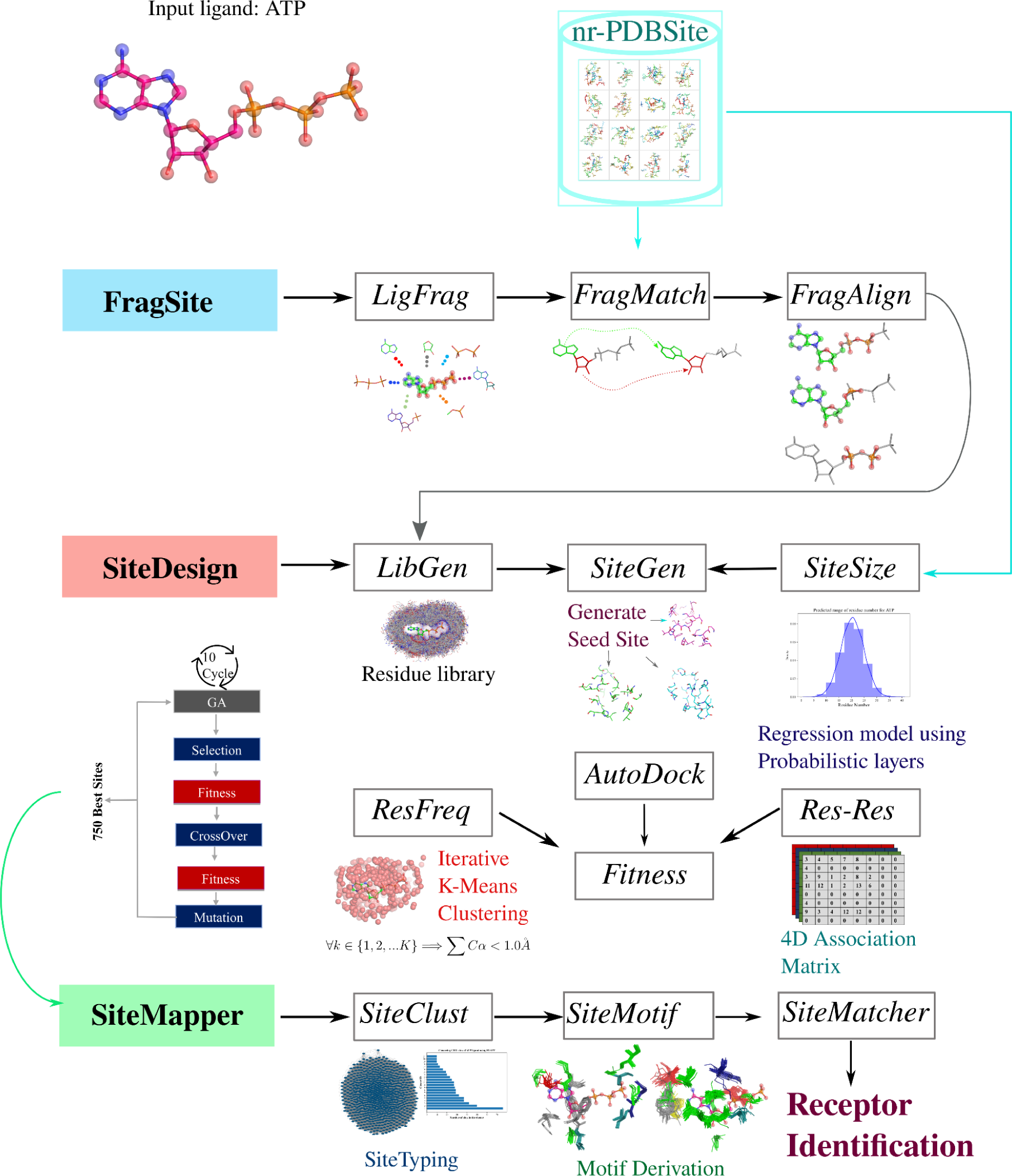
The architecture of CRD illustrates three principal modules, **FragSite**, **SiteDesign** and **SiteMapper**, each containing several functions. **FragSite** has the functions *LigFrag* that generates ligand fragments, *FragMatch* that finds common fragments in comparable ligands from PDB and *FragAlign* which aligns the matched fragments onto the query ligand. The **SiteDesign** module takes the FragAlign output and designs sites for the query ligand with the help of functions *LibGen*, *SiteGen*, *Fitness*and *SiteSize*. *LibGen* generates a residue library starting from the *FragAlign* output, while *SiteSize* determines the optimal residue range for the query ligand, outputs from both informing the *SiteGen* function which is based on a Genetic Algorithm that designs a set of possible binding sites for the ligand. The *Fitness* function has 3 components generated by subfunctions Autodock, Res-Res and ResFeq that help in locating dense residue regions and determining residue-residue associations and together guide the design by the *SiteGen*. The **SiteMapper** module post-processes the designed sites by clustering them into site-types by using Louvain, generating 3D site-motifs using *SiteMotif*, and finally using *SiteMapper* to identify possible receptors for a given ligand.

The first module of CRD that we develop is for ligand fragmentation, finding similar fragments in PDB and aligning the fragments to generate fragment sites, which we refer to as ‘***FragSite***’. We define a fragment as a set of connected atoms of length greater than four, with each atom making at least one covalent bond with another atom of the fragment. FragSite has 3 functions, *LigFrag, FragMatch* and *FragAlign*. *LigFrag* generates an exhaustive set of overlapping fragments for each ligand and internally uses CheckMol to ensure that known functional groups are not fragmented [17]. *FragMatch* compares the generated fragments against all PDB-ligands through a systematic fragment-vs-ligand similarity search, using a graph algorithm (Fig. S1). For this, each ligand in PDB is represented as a distance matrix (D) with atoms constituting nodes and edges drawn between them representing their Euclidean distances, with 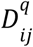 representing the distance between each atom ‘i’ to all other atoms ‘j’ of the query ligand ‘q’. As the clique detection involves searching all possible common paths between graphs, this is regarded as a combination problem and is often computationally intensive. To accelerate the execution, we implemented blazing-fast state-of-the-art algorithms used to detect binding site similarities at large scale (similar to that in [18]). We look for all possible matching cliques in the graphs of length greater than four. From the distance matrix (D), pairs of connected atoms are identified from the 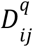 whose distance difference is less than 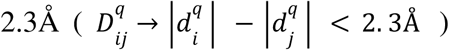. The cutoff 2.3Å imposes that the picked atom pair is covalently linked which will serve as the starting seed pair. The chosen seed pair is matched against distance matrices of all ligands in PDB 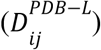. The criteria for matching are that (i) the differences between two seed pairs is 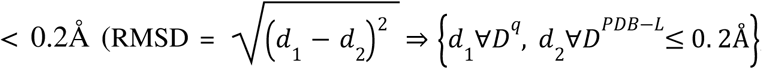, *where* 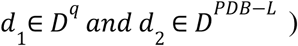, and (ii) the atom types of comparable pairs are similar. To prevent large memory access on CPU, only the top 20 percentile of matches ranked by matched fragment length are taken for subsequent analysis (Supplementary-Text-1). The *FragAlign* function superposes each of the top-matched fragments onto the query ligand using the Kabsch least squares-fit algorithm [19]. The RMSD of fragment alignment was observed to be < 0.2Å for most cases (Fig. S12).

The second module of CRD **SiteDesign has 4 functions-*LibGen*:** that constructs a spatial residue library, ***SiteGen***: that generates candidate sites, ***Fitness***: that evaluates and selects the fittest sites. Further, we develop an independent function ***SiteSize:*** that SiteGen consults, which defines the size range of the site for the query ligand. The Fitness function has additional sub-functions *ResFreq*: that constructs a residue conservation score and *Res-Res*: that constructs a residue-residue co-occurrence matrix. In ***LibGen,*** for each fragment in 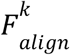, it’s binding site is extracted from the parent protein-ligand complex in PDB, and placed in the superposed framework of the query ligand, by applying the superposition matrix that was applied to align the matched fragment to the query ligand. This yields a large pool of amino acids transferred to the common frame of the query ligand, which in essence is a spatial site residue library specific to the query ligand, which we refer to as the ‘**Residue library**’ (Fig. S2 showing examples for four ligands - ATP, GLC, SAM and FAD).

The residue library is very dense and can lead to the generation of an innumerable number of residue arrangements, from which the most likely ones are selected. **SiteDesign** uses a Genetic Algorithm and selects feasible arrangements, and outputs a large number of candidate sites. ‘SiteGen’ takes in the fragsites output from the first module and designs optimal sites. A detailed explanation of the SiteGen module is given in supplementary-text-2 (Fig. S11). As a recursive algorithm, SiteGen is repeated multiple times until the desired number of residues is obtained. Given the coordinate space of ligand L, we strive to maximize the search space region S that will hold the optimal number of residues, which prevents over-housing.

The challenge however, is in defining an optimal size of the site or in other words the number of residues in the site for a given ligand. To address this, we develop ‘***SiteSize***’ function, which combines a structural bioinformatics analysis and a machine learning algorithm to learn patterns from known structural data. A given ligand is typically seen to be housed in a site of a similar size irrespective of the size or nature of the protein in which it occurs (eg., ATP binding sites are in the range of 19 to 28 residues). To factor this type of knowledge systematically for any given ligand, especially when the structure of the query ligand or its analogue is not available in PDB, we define the site range in terms of the *MinBound* and *MaxBound* values which represent the minimum and maximum number of residues required to construct the binding site. This serves as the boundary conditions within which the search space ([MinBound, MaxBound]) must be explored in order to generate residues. We define a new objective function (MinMax.py) which takes in an input of physicochemical descriptors of ligands in PDB using the PyBioMed package [20]. A total of 456 features 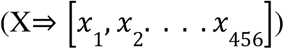 representing topology, connectivity, and QSAR properties for each atom in each ligand is computed. The predictor variable (output) is the number of residues encompassing the binding site. Using tensorflow probability, we developed a probabilistic regression method based on a deep learning model (f(y) ⇒ 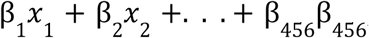, *where* β *is a statistical parameter*). The advantage of probabilistic regression over traditional regression is that instead of predicting one single numerical value, we can estimate prediction intervals drawn from the distribution. We used a loss function of negative log likelihood (-log P(y|X)) which was minimized by *Adam optimizer* at a default learning rate of 0.01.

In the ***Fitness*** function, we developed a ‘triplet fitness’ score that evaluates candidate sites generated by SiteGen based on (a) interaction-energies, residue preference in the designed elite members and (c) extent of residue-residue co-occurrences in pairs, triplets, quadruplets and so on (Fig. S3). The triplet fitness score is aimed at preventing overfitting due to electrostatic potential optimization and eliminating putative sites that have unfavorable interactions, poor atom connectivity or poor geometries (Supplementary-Text-3). We combine AutoDock inter-molecular energy scoring function with two additional fitness functions i) Residue conservation fitness (*ResFreq*) and ii) Residue-residue co-occurrence based association (*Res-Res*). Combining the three scoring terms allows us to search for a large number of potential residue combinations without creating an overfit.

AutoDock scoring function is an empirical free energy scoring function that estimates binding energy of a small molecule ligand to a protein, based on the Amber force field and consists of pairwise terms for dispersion/ repulsion, hydrogen bonding, electrostatics, desolvation and loss in conformational entropy upon binding of a ligand to a protein (TotalEnergy = ^E^_bond-angle_^+E^_bond-length_^+E^_dihedral_^)^ [21].

The function ‘***ResFreq***’ determines the frequency of occurrence in the residue library. It starts by identifying highly occupied regions around the query ligand’s fragments, which is achieved by an iterative k-means clustering algorithm. Clustering is attempted systematically at increasing k-values from which we choose the optimal number of clusters, based on an elbow-plot.

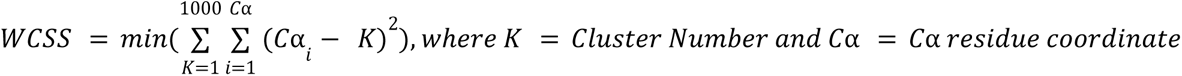

This yielded spatial positions that were densely occupied by residues around the ligand (Fig. S2). The number of residues whose Cα is within 1.0Å of the cluster centroid (Cn) was taken as the degree of conservation of that cluster. This denotes the conservation fitness terms.

From the Residue Library, we compute a 4D matrix by scanning the library to compute the residue types, frequency of each of the 20 amino acid residues in the vicinity of each ligand atom. The fitness function used in the ***SiteDesign***, consults the 4D matrix for evaluation of each candidate site. The frequencies of all the 20 amino acid types in a given cluster is captured in the 4D matrix of shape KxLxMxN, where K represents number of PDB matched fragments and L equals number of ligand atoms. The *Res-Res* function scans the clusters and derives residue-residue spatial co-occurrences as pairs, triplets, quadruplets and so on. This is accomplished by calculating the distance of each residue from all atoms of ligand (L) and storing them in the MxN matrix, where M represents 20 amino acids and N stores the actual distance.

During initialization, N will be an empty array of length 10, set to zero. For each ligand atom, the distance of each residue is encoded sequentially to N. This 4D matrix will be examined each time a new site is constructed to determine how frequently the residue-residue close proximity is detected.

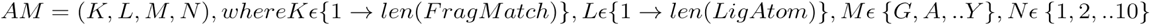

For each new site candidate, the 4D matrix will be consulted to check how many times the residue-residue nearness is spotted. The use of the *Res-Res* association matrix together with *ResFreq* residue conservation will help us identify residues that are frequently present and co-occur in pairs, triplets or in any higher order.

Evaluation of the Triplet fitness function: To test the efficiency of triplet function over pure energy-based methods such as AutoDock, 100 sites were randomly generated using SiteGen and ten best sites were chosen independently based on (a) AutoDock function alone and (b) the Triplet function. This was repeated in 10 different runs, resulting in 100 best generated sites. 100 sites generated for each of the four ligands were compared with the corresponding known PDB sites using FLAPP which was then quantified for similarity with F_min_ > 0.4 (Fig. S4). From Fig. S4, it was evident that the best sites selected from Triplet function fare better in mapping to the most of the known site complexes as compared to AutoDock alone. Next, we evaluated the importance of each of the three functional terms in the triplet fitness function using the boruta algorithm, which is based on a random-forest classifier and widely used for feature selection [22]. We observed that all three functional terms were identified to be the top-ranked feature in selecting the best matching sites in one or more of the tested cases (Fig. S5) indicating the importance of the combination and hence the rank of all three was used to quantify the fitness of the generated sites.

The SiteGen module outputs 100 fittest candidate sites in each generation and by default, runs for 5 generations. 15 elite members are taken from each generation leading to a total of 75 sites per run. 10 such runs are performed, which yields a total of 750 designed candidate sites. The fourth module, ‘***SiteMapper***’, which is a post-design analysis module clusters designed sites, finds cluster representatives, matches them with known sites in PDB, selects key residues in the sites and predicts the closest cognate receptors for a given ligand from PDB. To achieve this, SiteMapper uses 3 functions, *SiteClust*, which clusters the 750 best designed sites to identify similarities among themselves and identify site-types and *SiteMotif*, which derives structural site motifs and *SiteMatcher*, which compares the motifs against a comprehensive PDB-derived sites database (NRSiteDB). SiteClust carries out an all-vs-all similarity of the 750 sites using FLAPP, a recently developed in-house fast-matching algorithm and identifies site-types by clustering them using the Louvain algorithm. Each cluster is regarded as a site-type and for each site-type 3D site motifs are generated using SiteMotif [23]. Cluster membership is given when each member has a RMSD < 1Å with majority other members), 3D site motifs capture the most frequently occurring site residues in the clusters. The clusters are ranked by their size. *SiteMatcher* takes in the designed sites as an input and compares them against NRSiteDB, for identifying matching binding sites (F_min_ > 0.7 or no. of alignment; n > 12). The proteins housing the matched sites are predicted as receptors for that ligand. A docking run is then performed to estimate the theoretical feasibility of binding of the ligand to the predicted receptors.

### CRD reliably generates accurate sites for different ligands

We evaluated the performance of CRD to predict receptors for well known ligands by testing whether it can design known binding sites across four distinct ligands. We selected these ligands from a systematic analysis of projecting all small-molecule ligands bound to proteins available in PDB (PDB-ligands) into a 2D matrix with partition coefficient (logP) and molecular weight as the two axes and selecting ligands from different densely populated locations in the 2D matrix (Fig. S6). The ligands ATP, SAM, FAD and Glucose were selected which are seen in several protein-ligand complexes in PDB and are recognized by proteins belonging to diverse sequence families and perform diverse functions. The proteins that bind to these are known to exhibit large structural diversity with multiple structural folds not only across the ligands but also within each ligand [24,25]. Put together, this represents a challenging design space and hence suitable to evaluate the performance of a multi-module multi-step method for predicting receptors, given a query ligand.

We collected a dataset of 55,301 protein-ligand complexes (NRSiteDB) for these four ligands from PDB and excluded their entries from the NRSiteDB. After ensuring that all instances of these ligands in PDB are excluded for use in the Fragsite module, we ran CRD and tested if it was capable of designing sites that mimicked natural sites. For this, we checked using FLAPP, if the designed sites shared similarities with one or more of the known sites (F_min_ >0.4, no. of alignment n > 6). Greater the F_min_ score, higher is the similarity with the known sites. The search space for our designed sites spans the entire landscape of theoretically feasible site architectures for binding a given ligand and we verified that all ligand atoms and the site space around the ligand atoms were well represented in the FragAlign and LibGen respectively (Fig. S2).

#### ATP

ATP is a large ligand with 31 atoms and containing a negatively charged phosphate end. For ATP as the query ligand, the FragMatch module found 1,17,737 PDB-ligand fragments that were aligned to the query ligand. The fragments were ranked by length of alignment and the top-ranked fragments were found to be identical to near-identical to the query. Since the query ligand in PDB is removed, FragMatch identified many ligands that are similar to the query (ADP, AGS and AMP). FragMatch also identified many similar fragments from otherwise dissimilar ligands (Tanimoto Coefficient < 0.3) bearing functional moieties similar to those in the ATP (Sheet1-FragmentsObtained.xlsx). Identification of a large number of fragments even when the ligand is removed in fact highlights the usefulness of the fragment-based approach. Different fragments of the ligand generated differently populated fragsites. The imbalance was addressed by taking the top 20 percentile of matches for each query fragment, yielding 10,808 best fragment matches, from which corresponding binding site residues from the matched proteins were extracted and aligned, generating a ‘residue library’ of 34,838 amino acids (Fig. 2b). The Res-Res module was then applied onto the ‘residue library’ yielding 3735 k-means centroids weighted as the number of residues with 1Å of each centroid. SiteSize identified site sizes for ATP to be in the range of 20 to 26 residues (Fig. S7a). SiteGen, coupled with the Fitness module, outputs 750 top-ranked designed sites for ATP recognition. The SiteClust function in SiteMapper module identifies that the designed sites belong to 14 clusters, each representing a distinct enough site-type (Fig. 2c). 10 out of 14 motifs showed matches (F_min_ >0.4) against known ATP sites, indicating the CRD design to be successful in mimicking known sites (Fig. 2e). 630 known sites for ATP are present in PDB, which when compared amongst themselves in an all-vs-all comparison using SiteMotif, identifies 15 clusters. 7 out of 14 designed sites types match with 7 of the known site clusters (Fig. 2c). This clearly shows that CRD successfully designed multiple known site-types for ATP. Next, the SiteMatcher function in SiteMapper carried out an exhaustive comparison of the 750 designed sites against 55,301 sites in NRSiteDB and the proteins housing these matched sites are identified as predicted receptors for ATP. The top-ranked receptors showed very high similarity with known ATP binding proteins (F_min_ > 0.7), (top 2 being proteins pseudokinase and CDK2 (Fig. 2f)). A docking run is performed to estimate the theoretical feasibility of ATP binding to all the identified hits from NRSiteDB, which indicated the intermolecular energies to be favorable (Sheet-1 ReceptorDocked.xlsx).

**Fig. 2.**
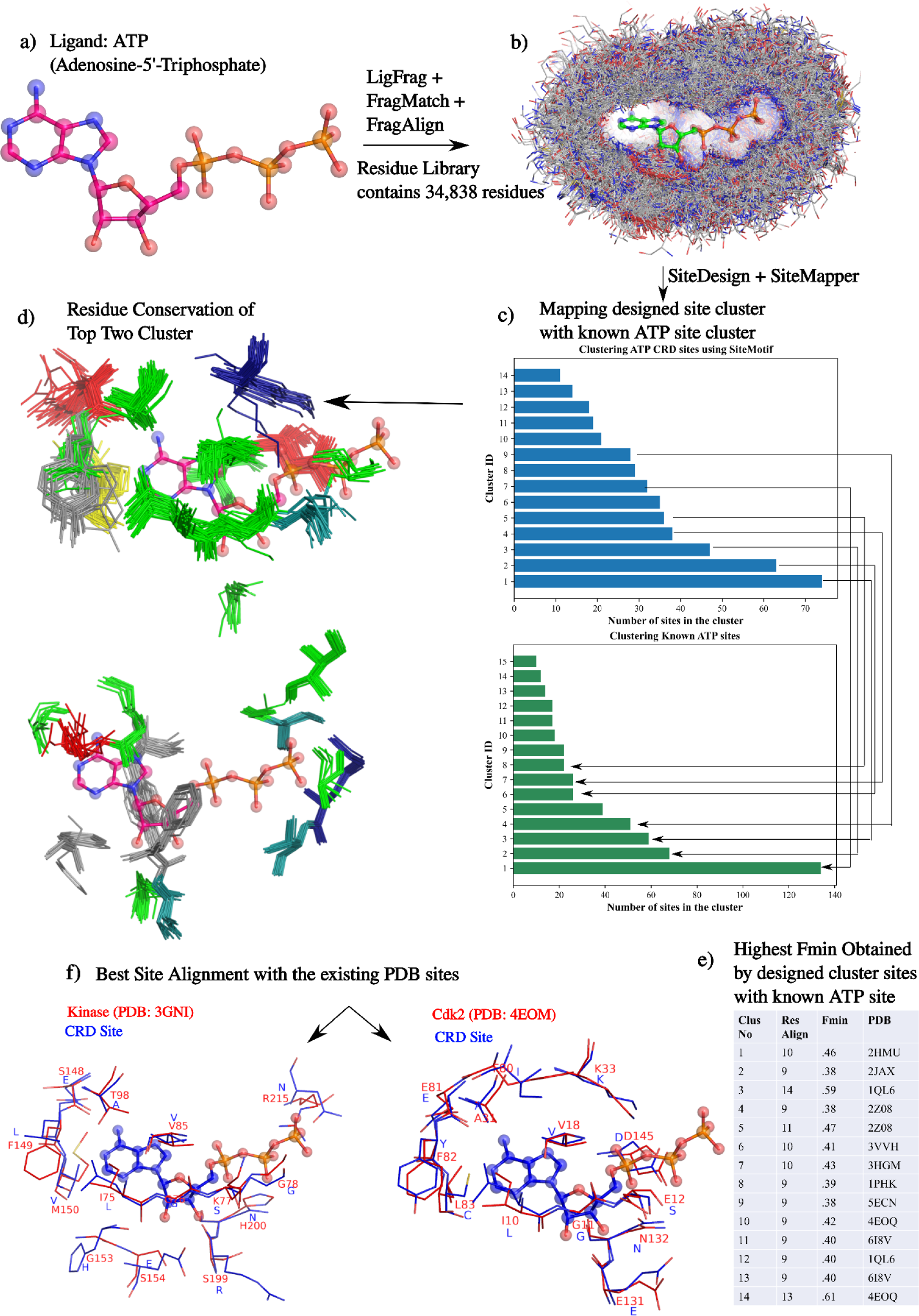
Design of binding sites and prediction of receptors for ATP. (a) Starting from coordinates of ATP in PDB format as input, (b) the FragMatch functions: LigFrag, FragMatch, and FragAlign were executed sequentially. 34,838 residues formed the Residue Library that completely encircled the ligand atoms, (c) using which SiteGen in the SiteDesign module generated 750 best scoring sites. d) SiteMotifs are constructed for the largest two clusters identified after comparing all 750 designed sites among themselves. Residues are coloured using a standard scheme (basic-blue, acidic-red, neutral-green, aromatic-grey, small uncharged-teal). A lysine and an aspartate are seen to be conserved near the phosphate moiety of ATP aromatic amino acids are conserved near the adenine ring in both clusters. (c) CRD site clusters were compared to the site clusters obtained from 638 known ATP-PDB site complexes, and 6 out of 14 designed site clusters matched the known ones. (e) The highest F_min_ obtained by each cluster member with the known sites are shown. (f) Receptor prediction by Sitemapper showing two highest scoring pairwise alignments (by comparing all 750 sites to 689 known ATP sites), as the ATP binding pseudokinase STRAD (PDB:3GNI) and ATP binding CDK2 (PDB:4EOM), Showing how closely CRD designed sites mimic the actual sites.

#### SAM

SAM is an uncharged ligand with 27 atoms, differing from ATP in having a methionine moiety instead of the tri-phosphate end. Taking all the known SAM sites from NRsiteDB and clustering them as in the case of ATP, indicated that there are 8 known site-types for SAM. For SAM as the query ligand, CRD designed sites of 8 types, matching with all the 8 known types. The top-matches were observed to be known SAM binding sites (top 2 receptors being arginine methyltransferase and catechol methyltransferase, Full analysis in Supplementary-Text-4, Fig. S8).

#### FAD

FAD is a large uncharged ligand with 53 atoms, a widely used redox-active coenzyme involved with several enzymatic reactions in metabolism. Taking all the known FAD sites from NRSiteDB and clustering them indicated that there are 14 known site-types for this ligand. CRD designed 14 site types, of which seven matched with the known FAD site types). The top-matches were observed to be known FAD binding sites (Fig. D9f) corresponding to proteins Renalase and Histone demethylase (Full analysis in Supplementary-Text-5, Fig. S9).

#### Glucose

Glucose was chosen as a representative of small, highly polar ligands. Taking all the known glucose sites from NRSiteDB and clustering them indicated that there are 7 known site-types for this ligand. CRD identified 11 site-types for glucose, which corresponded to all of the 7 known site-types. The top-matching sites (Fig. S10f) and their associated receptors are identified as glucokinase and hexokinase, which are known glucose binding proteins (Supplementary-Text-6, Fig. S10).

### CRD generates similar sites for Similar ligands - yet distinguishes among them

We next asked if CRD designs similar sites for similar ligands and whether it has sufficient resolution power to discriminate between similar ligands. We chose a set of highly similar ligands - Glucose, Galactose and Mannose which differ only in the stereochemical orientation of one of their hydroxyl groups (Fig. 3a). This set was chosen to address a challenging problem to discriminate among their sites. Glucose has an axial O2 whereas it is at an equatorial position in mannose. Mannose is a C2 epimer of glucose. Likewise glucose and galactose differ only in the O4. Galactose is a C4 epimer of glucose. The networks of the 750 sites generated for each ligand were combined into a mega-network by drawing edges between all pairs of sites that were similar (F_min_ > 0.5), (Fig. 3b), which clearly shows that they form dense connections among themselves but also overlap extensively. Clustering of the mega-network indicated 17 clusters with sites designed for mannose, glucose and galactose predominant in clusters 2, 4 and 1 respectively, though interspersed among others, clearly demonstrating that the CRD designed similar sites for similar ligands, but at the same time has the resolution power to differentiate among them. 397 complexes of glucose, mannose and galactose are available in NRSiteDB. We extracted these and compared all-vs-all among themselves and found that there was overlap in many of the members from one ligand with that of the other (Fig. 3c). CRD designed sites indeed followed the same trend (Fig. 3c). To analyze the differences among the sites for the 3 ligands, we compared them at a higher threshold of similarity (Fig. 3e). We selected clusters from the network that shared the highest similarities between ligands and observed that the glucose cluster has prominent asparagine residues juxtaposed to interact with the O2 hydroxyl. The mannose sites have an aspartic acid in that region. Likewise the mannose designed sites differ from that of galactose in having an aspartate residue interacting with the O5 instead of an asparagine or alanine present in the galactose cluster.

**Fig. 3.**
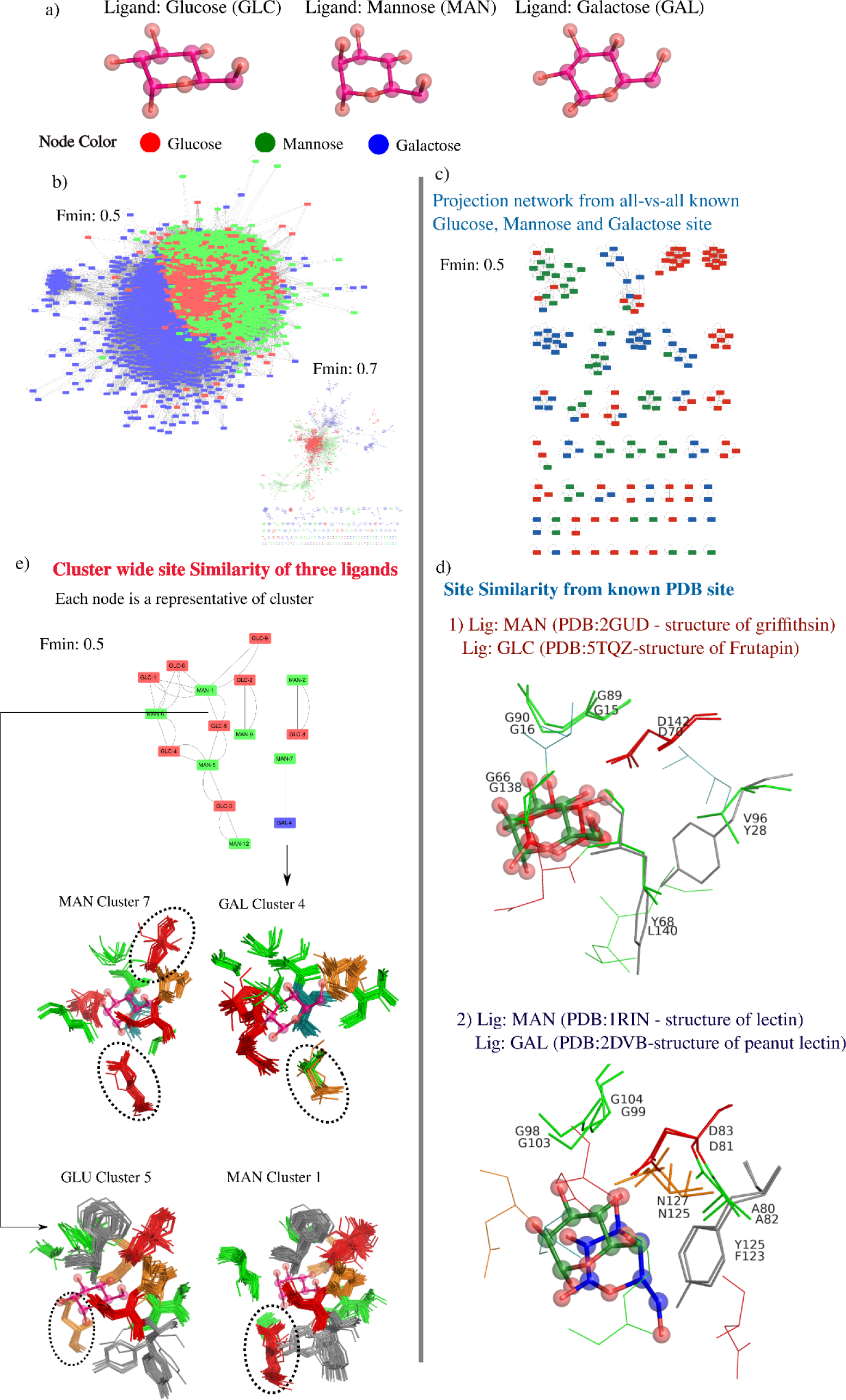
Testing CRD resolution to distinguish sites of closely related ligands. a) glucose, mannose, and galactose, b) An all-vs-all site similarity network of the CRD designed sites for the three ligands (red-glucose, green-mannose, blue-galactose) where nodes are the designed sites and edges indicate similarities between sites (F_min_ > 0.5). The network was constructed by grouping all three sets of 750 sites (750*750*750 = ∼0.42 billion pairwise combinations). High extent of similarities among them yielded one large mega-network. The network also shows that the binding sites of these 3 ligands tend to form denser connections among themselves (seen more clearly at a higher similarity threshold network of Fmin 0.7, shown as an inset) indicating that they are to some extent distinctive from each other. (c)A similar analysis was conducted on the binding sites of known sites from actual sugar binding proteins (GLC, MAN, and GAL), showing the same trend, further illustrated in (d) showing pairwise site alignments of PDB sites of MAN-GLC and MAN-GAL site combinations. (e) The same network of CRD designed sites as in (b) but each node representing a cluster of sites of glucose (red), mannose (green) or galactose (blue), from which Site motifs (standard residue coloring) are generated and compared: Both Mannose and Glucose (1 and 5) clusters were highly similar with differences occurring mainly in O2 region, where aspartate was seen to be conserved in the mannose site, whereas asparagine was found to be conserved in the glucose site. Similarly, cluster-7 of mannose sites shared similarity with cluster-4 of galactose sites with a difference being the presence of more aspartates in the mannose site as compared to the galactose site.

### CRD predicts receptor classes for several drugs

Next, we investigated how well CRD fares in designing sites for drug compounds. We chose two drug families (Penicillin G, Ampicillin, Amoxicillin and Methicillin) and non-steroidal anti-inflammatory agents (NSAIDs - Ibuprofen, Naproxen and Celecoxib), and prepared the fragset after removing any instances of them in PDB. CRD designed 9 site-types for Penicillin G, 2 of which matched closely with the known sites of penicillin binding protein and beta-lactamase respectively (Fig. 4). In addition, Guanylate binding protein, Monoamine oxidase and Histone demethylase are predicted to be possible receptors by CRD. Docking of these ligands to these proteins indicated a mean binding energy of -10.1, -9.1, and -9.0 kcal/mol respectively (better than the binding energies with penicillin binding protein with a mean binding energy of -6.7 kcal/mol), implying that theoretically binding would be favorable (Table. S1). Similarly, CRD predicted penicillin-binding protein and beta-lactamase among the top-ranked receptors for ampicillin, amoxicillin, and methicillin compounds as well (Fig. 4). With NSAIDs - CRD designed 21 clusters for ibuprofen, 5 of which matched with the sites of known complexes of ibuprofen which are with Cyclooxygenase-1 (COX-1) and Cyclooxygenase-2 (COX-2) proteins. Similarly, cyclooxygenases were among the top hits for naproxen and celecoxib (Fig. 5). GTPase and Ras were predicted to be additional possible receptors. Docking of these ligands to these proteins revealed a mean binding energy of -9.6 kcal/mol and -9.4 kcal/mol respectively (better than the binding energies with COX with a mean binding energy of -7-7 kcal/mol), implying that binding would be favourable (Table. S2).

**Fig. 4.**
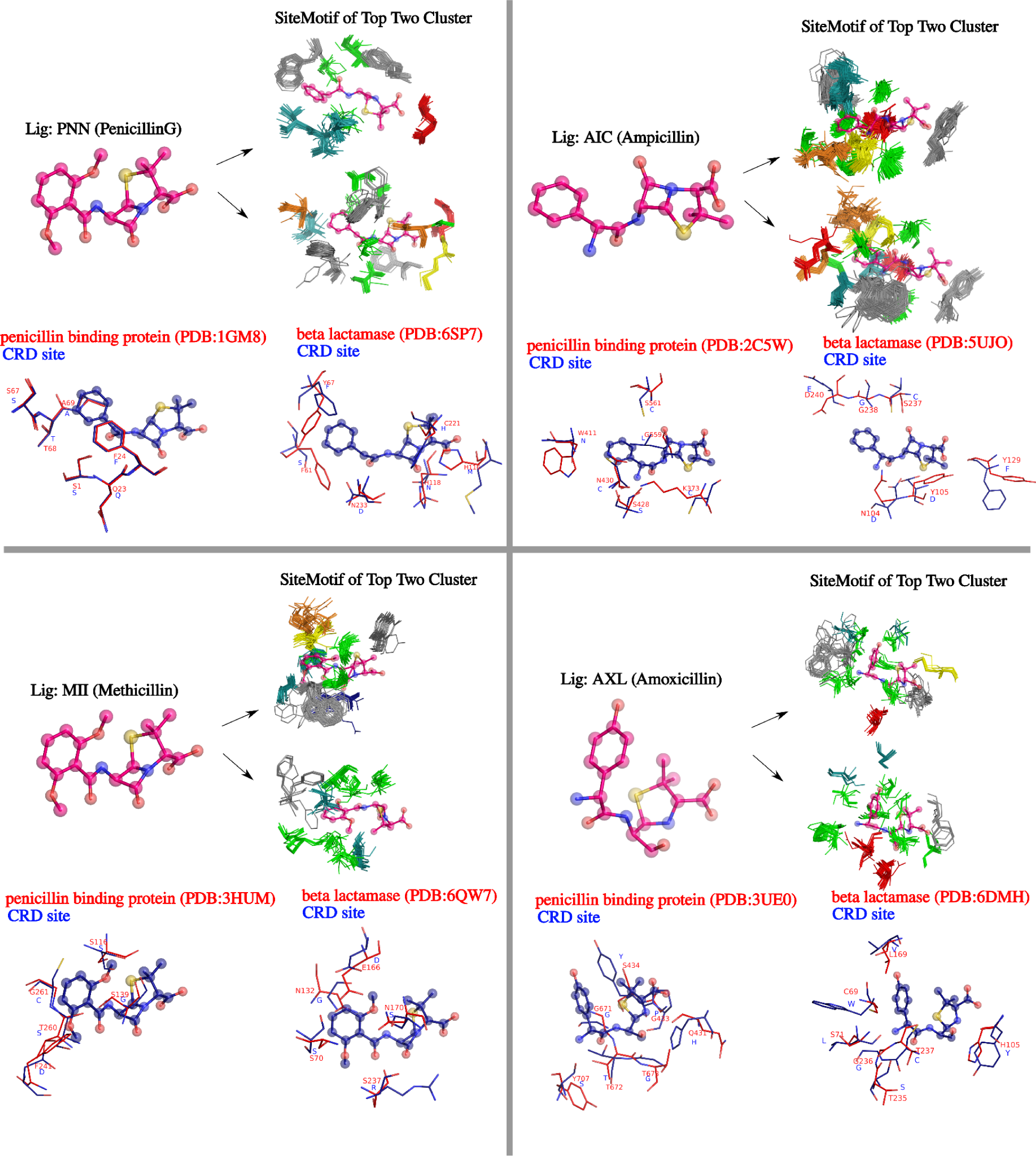
Examining sites designed for the penicillin family of drugs. Here, we have taken 4 widely prescribed penicillin drugs namely: PenicillinG, Ampicillin, Methicillin and Amoxicillin, and derived site types for each separately. The designed sites for each drug were compared with NRSiteDB, and it was found that CRD was able to generate a site with good residue-residue alignment with the known interaction partners (penicillin binding proteins and beta-lactamase). On the basis of these findings, CRD has the potential to design a binding site that mimics the binding site of actual drug binding proteins.

**Fig. 5.**
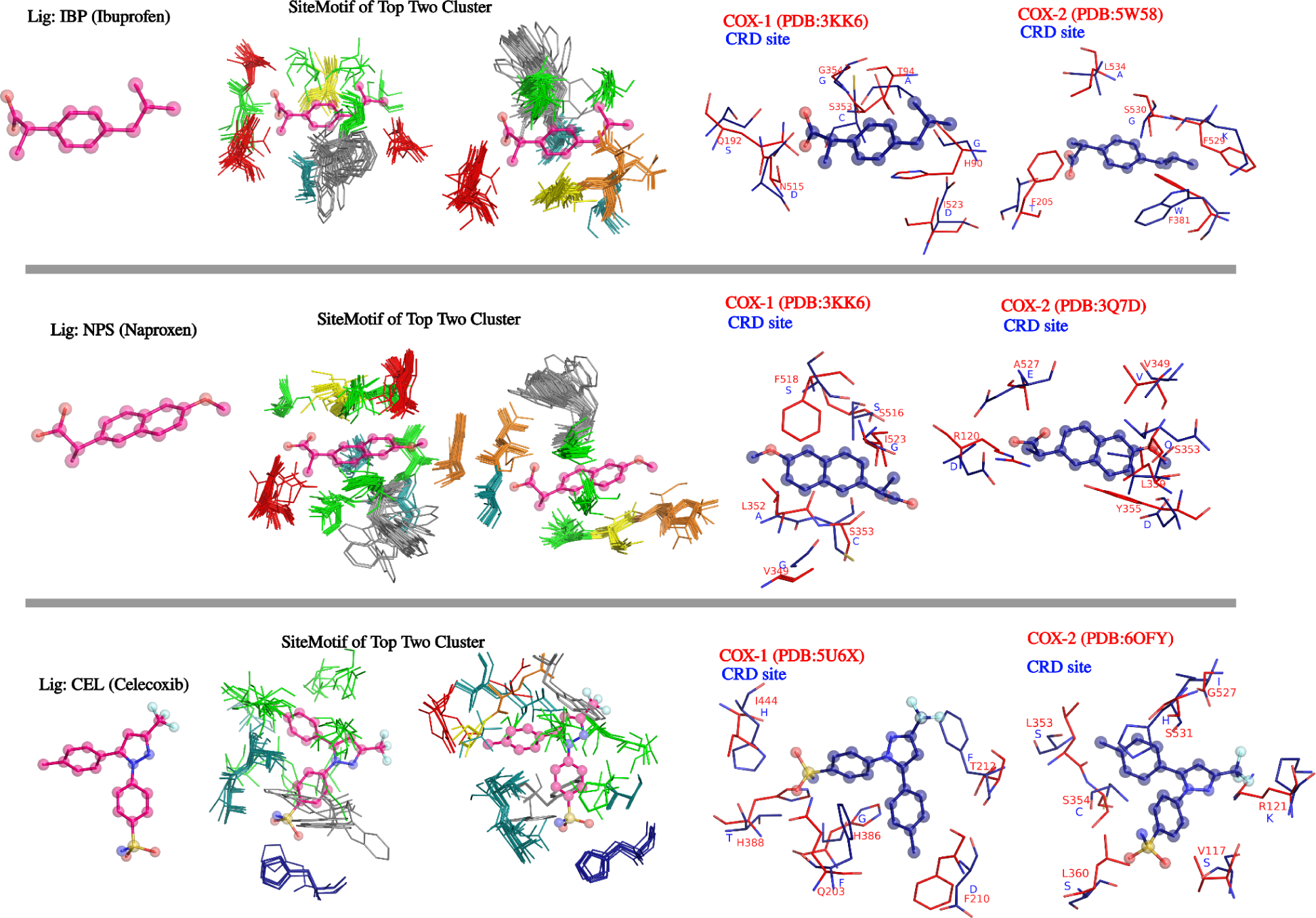
CRD sites designed for NSAIDs (Ibuprofen, Naproxen and Celecoxib) with site motifs for the top 2 clusters. SiteMapper identified high scoring matches with actual sites in cyclooxygenases (Cox-1 and Cox-2) which share high similarity with each other. CRD identified the broad receptor type but cannot predict selectivity.

## Discussion

Much of present-day biology relies on finding similarities among molecules at the sequence, fold, structure and expression levels and using them for drawing on the functional knowledge. As several biochemical processes are driven by specific sets of protein-small molecule interactions, binding sites in protein structures provide a rich source of information that can be leveraged to gain fundamental insights into protein function and also in drug discovery. Understanding how proteins and ligands recognize each other has remained a topic of great interest, leading to the development of many tools for predicting binding sites (Fpocket, PocketDepth, etc), comparing them and applying them for applications such as lead candidate design and drug repurposing. Currently, while there are computational methods that enable the prediction of a small molecule ligand for a given protein, predicting a receptor for a given ligand remains to be a huge challenge. In recent years, there have been efforts to utilize structural information to compare binding sites and extrapolate ligand-protein associations from known complexes to unknown protein or ligands based on binding site similarities. However these are also limited by being dependent upon known protein-ligand complexes and cannot be applied generally for any ligand.

Currently known ligand-protein associations are believed to be a small fraction of the actual associations, which means that there are a number of cases of protein-ligand complexes whose function and mode of achieving that function are poorly understood. This is especially true for allosteric regulators on one hand and understanding the effect of small molecule drugs on the other. A major limitation of phenotypically discovered drugs is the lack of understanding of the specific protein(s) that get targeted by the drug. Very often, one or two proteins that are modulated by a drug are identified which remain the principal focus for studying its pharmacology, leaving behind several other proteins that may actually play a major role. Identifying the cognate receptor(s) of a drug is a necessary step in understanding its mechanism of action and its off-target effects. Knowledge of the possible receptors of a drug is a very useful step in exploring its repurposing to another therapeutic indication. A general method to identify a receptor for any given ligand would not only throw light on the basic functional role of a given ligand or a receptor but could serve as a critical first step to address these and a variety of related questions. CRD bridges this key gap and provides a capability to predict receptor(s) at best and broad receptor classes as a minimum, for a given ligand.

CRD uses a de novo design approach to design putative binding sites and find proteins that may contain such sites. CRD is a suite of programs that integrates the following: (a) bio-chemo informatics based ligand fragmentation, fragment matching and subsite finding, (b) 3D graph-based methods for structural comparisons, c) site size estimation through machine learning, (d) a genetic algorithm for designing binding sites, (e) a physics-based triplet fitness function for evaluation and (f) site-mapping function for receptor identification. At each step, some of the key challenges have been addressed, making CRD capable of delivering the final predictions. Specifically, the fragment-based approach is a key enabling component of CRD, which ensures that CRD can cater to the design of any ligand, even when there is not even weak similarity to any ligand in PDB. This is because all ligands are broken down into chemical fragments which form the building blocks of organic small molecules. Fragmentation itself, if carried out conventionally, can be a restrictive step, as it typically generates known functional groups. In CRD, we overcome this challenge by exhaustively generating overlapping fragments and using each of them as an input to implicitly learn interaction environments. Residue placement and the residue selection are the next big challenges especially in terms of identifying the number and types in a given binding site. CRD precomputes the size range using a tensor flow algorithm, which goes a long way in deciding how many residues to place. The genetic algorithm based module selects residues such that the designed sites are the fittest. The choice of the fitness function is another critical parameter. Free energy estimations are routinely employed to measure strength of binding energy with ligands, typically used in docking simulations. We showed that the triplet function is superior to conventional energy-based function. Construction of a 4D matrix as a look-up table greatly facilitated computing the Res-Res, while K-means centroid as a look-up facilitates the Res-Feq components of the tripler fitness function. Analysis of the triplet fitness function also indicates relative importance of each feature, which can be used for ligand-specific adaptations to further prune the site-sets, in cases where that may be necessary. The next enabling feature of CRD is a fast comparison of the designed sites against the NRSiteDB, for which we used FLAPP, our recent superfast site comparison algorithm. The next feature of CRD is the use of clustering methods to group the designed sites and identify conserved residues within the clusters, which was enabled by SiteMotif, a recent development from our group. Put together, CRD is seen to be a powerful tool to design sites de novo and identify receptors for any given ligand.

We validated CRD by using a range of ligands and found that it could design sites very closely matching with known ones. We in fact ran CRD with and without retaining instances of the given ligand in PDB. In both cases we found that the designs were pretty successful. This clearly demonstrates the power of the fragment-based approach. For common ligands such as ATP, multiple known site-types are present in PDB, which are identified programmatically in our work, consolidating what is documented individually in literature. It was gratifying to observe that CRD was able to design these diverse site-types in a single run, indicating that it is able to capture different site arrangements that are feasible. Similar results were obtained with FAD, SAM and glucose, ligands with different properties. With the examples chosen for validation, it is clear that with similar ligands as query, similar sites are generated and yet it has the resolution power to distinguish among them.

CRD has some limitations as well. It outputs multiple 3D arrangements of residues which are grouped into different clusters, where sites within a cluster are seen to be very similar. From the examples tested, it appears that there is difficulty in ranking clusters merely based on size as the closest hit has in some cases been observed in lower-ranker clusters, thus making it difficult to output one best hit for the designed site. The clustering is critically dependent on the distribution of all-pair distances and hence the significance of the clusters are ligand-specific. In other words, 2 closely designed sites could be in the same cluster in some cases where as in different clusters in other cases. A further difficulty comes due to the challenges in measuring accuracy of the designed sites, one main reason being the difficulty in distinguishing situations where a designed site is ‘not seen naturally’ versus a designed site is possible naturally but ‘not seen in PDB’, as PDB itself is incomplete. Although the coverage of fragments in PDB is seen to be extensive, there is no uniformity in the frequency of occurrence of some ligands and hence some fragments. This bias indirectly cascades into the site design as more commonly seen residues have a higher chance of being selected in the fittest designs. A further minor limitation is that, as all fragments are superposed onto a given query ligand, the site design is partially biased to the conformation of the query ligand. This can however be overcome by running CRD for a second conformation of the ligand where required.

CRD is a novel algorithm generating a new capability of predicting a receptor for a given ligand. As structure prediction is advancing at a rapid pace with newer technologies including AlphaFold [26], the availability of protein structures is not likely to be a major bottleneck anymore. The critical need of the day is to make use of the structural information to understand function and to enable more and more accurate applications in drug discovery and other areas. We envisage CRD to be useful for fundamental rationalization of ligand binding capabilities required for understanding of protein function, understanding multiple sites types for a given ligand - deriving site equivalences and of course for several applications in drug discovery, such as finding targets for phenotypically discovered drugs, finding off-targets and designing drugs with higher selectivity. In drug discovery, the concept of pseudo receptor models has been used to generate presumed key interaction sites as anchor points to bind the given ligand, which provides valuable pointers for lead optimization or designing improved lead candidates. CRD, in some ways, can be regarded as a huge advance in this direction and is likely to bridge the gap between structure-guided design and ligand-guided design.

## Methods

### Data and Code availability

Structural data used in this work is from the publicly available PDB repository from the RCSB [2]. The CRD source code is written in python-3.9 of the anaconda distribution and comprises 15 python classes totalling 9500 lines of code. The program is hosted on GitHub (https://github.com/santhoshgits/SiteDesign) along with a YAML file for easy installation and execution irrespective of the OS platform and mitigates version conflicts that can arise due to usage of different python packages within the same script.

#### Receptor definition

In this work, we use the term ‘receptor’ in a general sense, and refer to all protein structures known to bind or predicted to bind small molecule ligands. The term ‘receptor’ therefore is not restricted to receptors in cell signaling pathways alone, but aims to include all protein classes such as enzymes, transcription factors etc.

#### Site definition

The term ‘binding site’ refers to a set of residues that form a pocket or a cavity in a protein that is capable of accommodating a small molecule ligand. We refer to the binding site in the PDB as a ‘binding site’ if it has been complexed with the given ligand, consistent with the widespread usage in literature. When proteins are complexed with ligands, residues whose one or more atoms lie within 4.5Å of any ligand atom constitute the binding site.

#### Fragment definition

CRD uses the idea of matching ligand fragments, a superset of functional groups, to generate binding sites. A fragment is defined as a set of connected atoms of length greater than four, with each atom making at least one covalent bond with another atom of the fragment. Fragments are extracted from ligand complexes present in PDB. A single fragment could contain multiple functional groups as long as they are covalently linked. Our notion of binding site design is that, if two comparable ligands share similar local substructure, such that any part of chemical spaces of query ligand overlaps with the atoms of PDB ligands, then the composition of residues around the vicinity of two comparable fragments will be similar [16,27]. This approach makes the analysis more methodical and meaningful as it allows us to discover large connected atoms present between two ligands. By default, only fragments having at least 4 atoms were considered. The CheckMol program was used to prevent fragmentation that could occur within large functional groups such as benzene or phenol [17].

#### NRSiteDB

Starting from all protein-ligand complexes solved by X-ray crystallography with a resolution of 2.5Å or better, available in PDB (v 30 April 2019), redundancy was removed by filtering based on sequence similarity at 70% sequence identity and removing duplicates, which resulted in a set of 55,301 protein-ligand complexes. Binding sites extracted from this set are referred to as NRSiteDB [2].

### Fragment Match

For every ligand in NRSiteDB, The FragMatch function finds alignable substructures common between itself and the query molecule. This is achieved by decomposing the 3D coordinate of ligands into 2D distance matrix and projecting each distance matrix as a clique detection problem. In graph terms, a distance matrix is nothing but an adjacency matrix which will enable us to create a graph data structure, with site residues as nodes and distance among them connected by edges. Since the number of traversals involved between two ligands are very large, the entire algorithm is written to support vectorization operation using LLVM compiler with additional support to parallelize on CPUs [28].

### Superposition of fragments and the site residues

Once the matching fragments are detected between the query and the target ligand from NRSiteDB, we extract residues present within 4Å of PDB fragments and transfer them around the corresponding atom of the input ligand via least square superposition using the Kabsch algorithm [19]. As the residues from several PDB entries will be transferred to one common frame, discrepancies such as missing atoms in a residue, unknown residue type, duplicate residue identifiers were addressed in LibGen. Residues of unknown type and those with missing atoms are excluded. Duplicate residue numbers were handled by renumbering them with unique identifiers in LibGen.

### Clustering of residues in ‘SiteDesign’ module

This step in the SiteDesign module identifies spaces with high probability of containing a binding site residue, ie., densely occupied with putative residues from the FragAlign function. This is required for avoiding overfitting in terms of adding too many residues in a site. An analysis of known binding sites by Baker and co-workers using RosettaHoles [28] has shown that binding sites of proteins are not densely packed. Several regions that are void, not occupied by any amino acid are evident in many structures. In the absence of the boundary condition, there is a danger of placing multiple residues that appear to be energetically advantageous, although not observed in nature. Clustering is performed by the k-means algorithm, optimal k decided based on the within-cluster sum of squares using an elbow plot. Clustering circumvents this step by eliminating all sparsely occupied regions for residue placement.

### The Search method to design binding sites

An innumerable number of sites can be made using the residues from the Residue-library. However, not every site made from sampling of the Residue-library can be regarded as a suitable binding site for the given ligand. Binding sites can have a number of unconnected amino acids, precluding the use of established rules such as peptide bonding or Ramachandran mapping which are routinely followed for protein design. Also, the use of standard energy functions such as potential energy calculated from AutoDock tends to overfit when applied on binding sites (Supplementary-Text-3). Given that the efficient placement of residues without the use of structural scaffold remains a difficult problem, we have devised a novel method SiteGen which is essentially a fragment-guided site builder, which leverages the Cα position of residues derived during fragment alignment step and residue conservation from the clustering exercise as a cue to build sites. A detailed explanation of the SiteGen module is given in Supplementary-Text-2. SiteGen is a recursive algorithm that is continued until the desired number of residues is found. Fig. S11 displays the SiteGen pseudocode.

### Evaluation of the fitness function

Using ‘Residue Library’ as the input, the SiteGen module generates binding sites conditional on the number of residues in the range predicted by the MinMax module. The suitability of the generated site was determined not based on traditional energy based function but via a custom objective function termed as the triplet function. In addition to calculating the potential energy, the triplet function also considers residues conserved around ligands and the associations between residues. We therefore evaluated the extent to which the triplet function identifies a true site as opposed to sites identified based on binding energies alone (Supplementary-Text-2, Supplementary-Text-3, Fig. S4, Fig. S5).

### Validation: Selection of ligands (2D matrix), ligand choice

A systematic exercise was carried out to select representative ligands for testing CRD. All PDB ligands were placed in a 2-dimensional matrix with molecular weight and partition coefficient (logP) as the 2 axes. Both molecular weight and LogP were calculated using open-babel. The 2D ligand matrix is shown in Fig. S6, from which densely occupied regions were considered (circled grids) and one representative ligand (the middle most member) in the grid was chosen.

### Execution of CRD and Parsing the outputs

To make our algorithmic suite easy to execute, three main wrapper Python scripts are included in CRD. They are PocketDesignInitiator.py, MinMax.py and PocketDesignGenerator.py. The program PocketDesignInitiator.py implements LigFrag, FragMatch, FragAlig, LibGen, Res-Res, ResFreg functions. Program MinMax.py executes the SiteSize function, which is independent of all other CRD functions. PocketDesignGenerator.py runs the SiteDesign module and outputs 750 sites. A number of ligands were tested using all three wrapper scripts. Keeping 4 CPU Cores and ATP.pdb as a query molecules, PocketDesignInitiator.py would take 4 hours to complete, MinMax.py would take 20 mins to complete, and PocketDesignGenerator.py would take ∼40 hours to complete.

By default, CRD(SiteDesign) was executed in 10 runs, with each run producing 5 generations of 100 individual sites in each, of which 15 best ones were selected as elite members. There is however a handle to change (a) the number of sites generated in each generation, (b) number of generations and (c) number of elite members that were carried forward from each generation and (d) number of runs (number of times the whole process is repeated). In addition, all the GA parameters such as the recombination and mutation frequencies can also be varied. The default run of CRD outputs 750 sites for each query ligand. These were compared all-vs-all using FLAPP with the following criteria: F_min_ > 0.4; no. of residues aligned n > 6. The all-pair distances obtained are used for clustering them using the Louvain algorithm [26]. Each cluster was regarded as a site-type, for which SiteMotif was used to identify 3D motifs conserved in that cluster. By default, each cluster has members that have similarities F_min_ > 0.7, among them and hence show deviation only in a few residues in the site. A representative set of residues such that it has the highest similarity to all other members in that cluster (by using a graph G: V (sites), E (similarity F_min_>0.4) and taking the highest degree vertex) was considered as the motif for searching against NRSiteDB. Alternatively all designed sites in the cluster can form the input.

## Data Availability

The source code of CRD is available at https://github.com/santhoshgits/SiteDesign. The NrSiteDB data needed to run it is available at https://zenodo.org/record/7780052.

## Supporting information

Penciliin-Class-Cognate-Receptors

ReceptorDocked

NSAID-Class-Cognate-Receptors

FragmentsObtained

## Supplementary Figures

**Figure S1.**
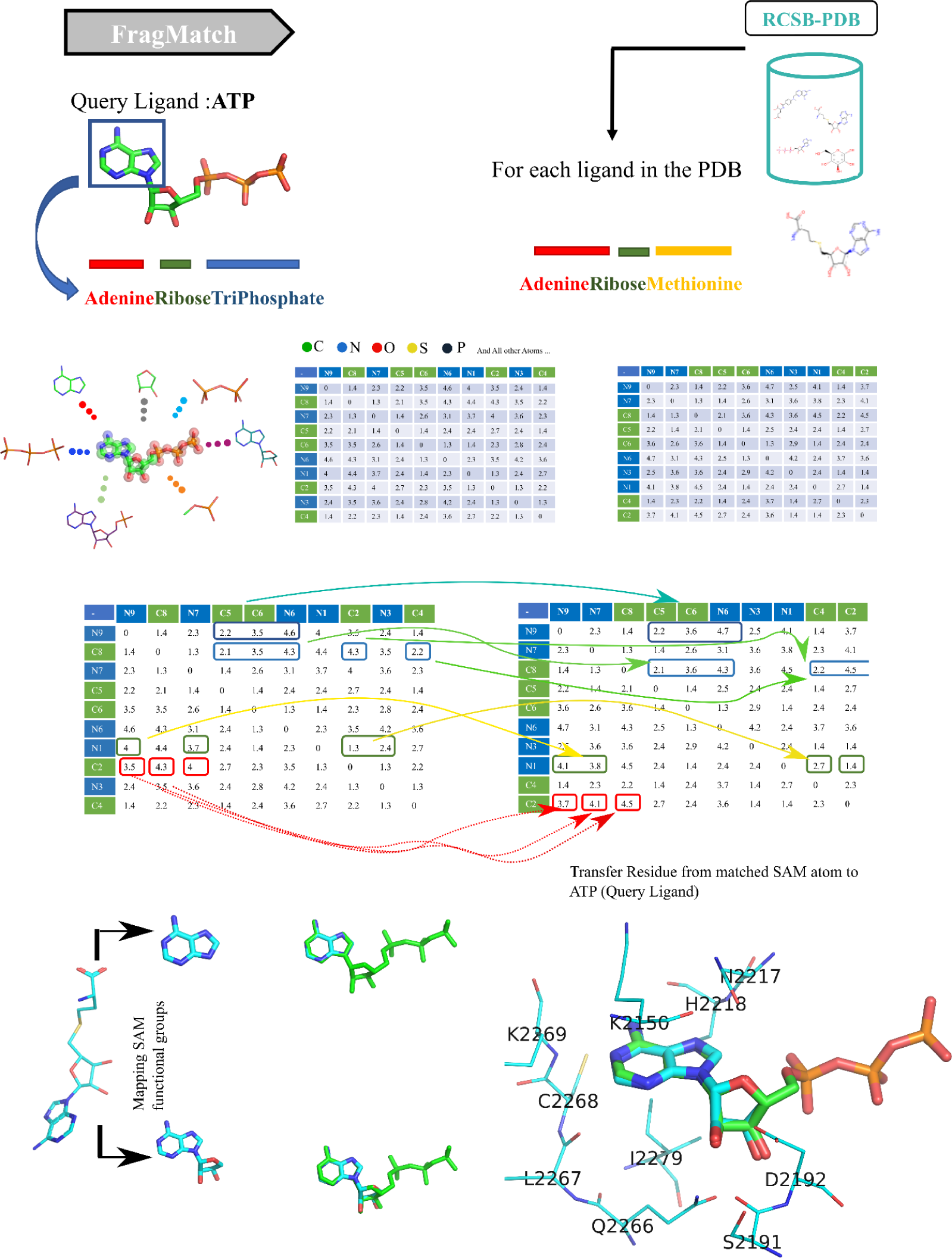
An illustration of N common substructure (N-CSS) implementation. An input ligand is decomposed into a 2D distance matrix of intra-atomic distances. A graph traversal search is then employed to look for the fragments that are common between the query molecule and the PDB ligand. Searches were optimized via use of vectorization and intel-SVML libraries for numerical calculations. Full alignment was obtained by applying the Kabsch algorithm to each of the obtained fragments.

**Figure S2.**
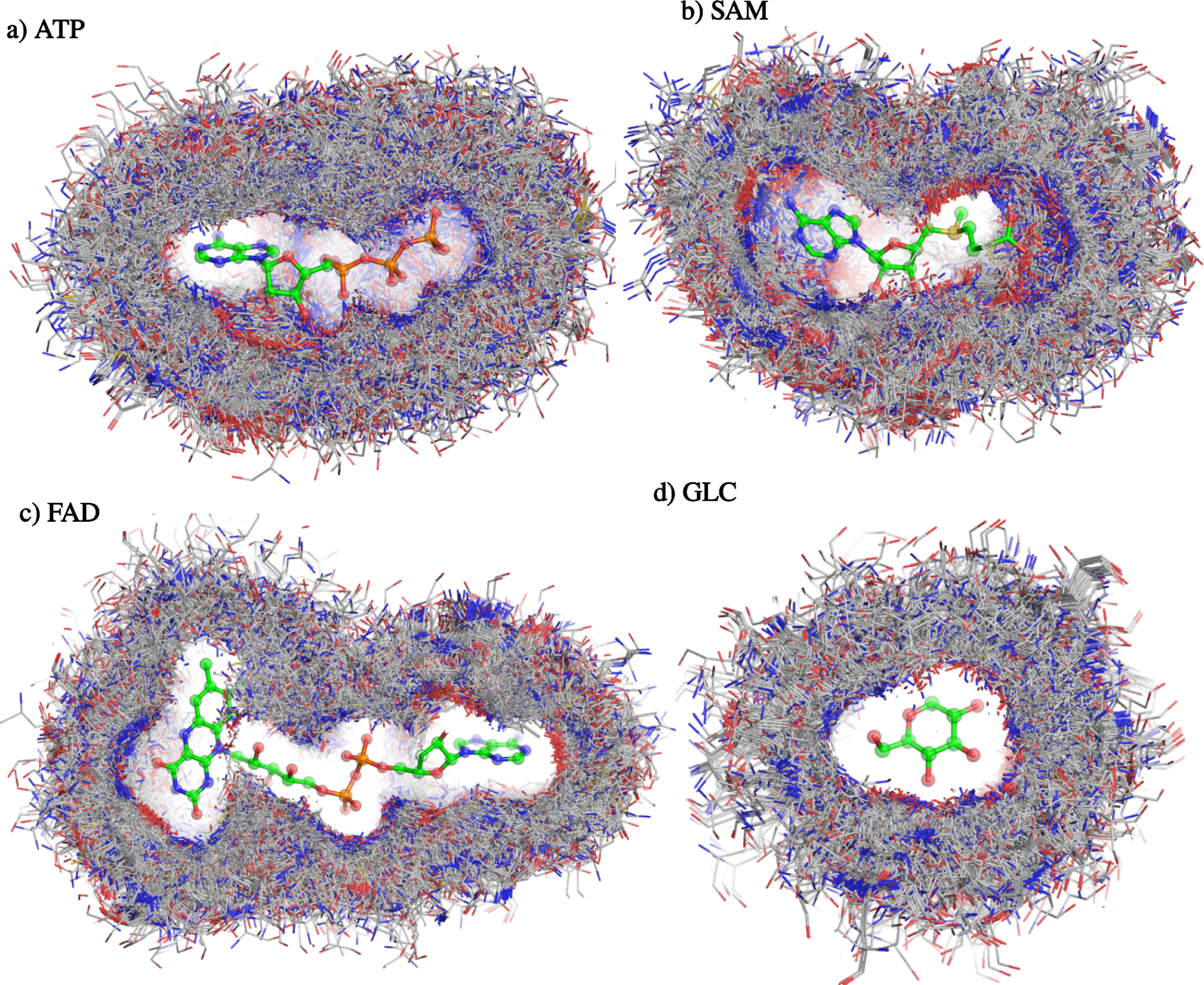
Generation of Residue-library using FragMatch and FragAlign. The FragMatch module was used to find matchable PDB ligands for each of four ligands namely: ATP(adenosine triphosphate), SAM(S-adenosylmethionine), GLC(Glucose) and FAD(Flavin-Adenine dinucleotide). FragAlign translates residues around PDB fragments to the query using the acquired atom-atom correspondence between query and PDB ligand. Applying FragAlign to each of matched hits generates overlapping residues that entirely occupy the surrounding volume of ligands, which is called ‘Residue-library’.

**Figure S3.**
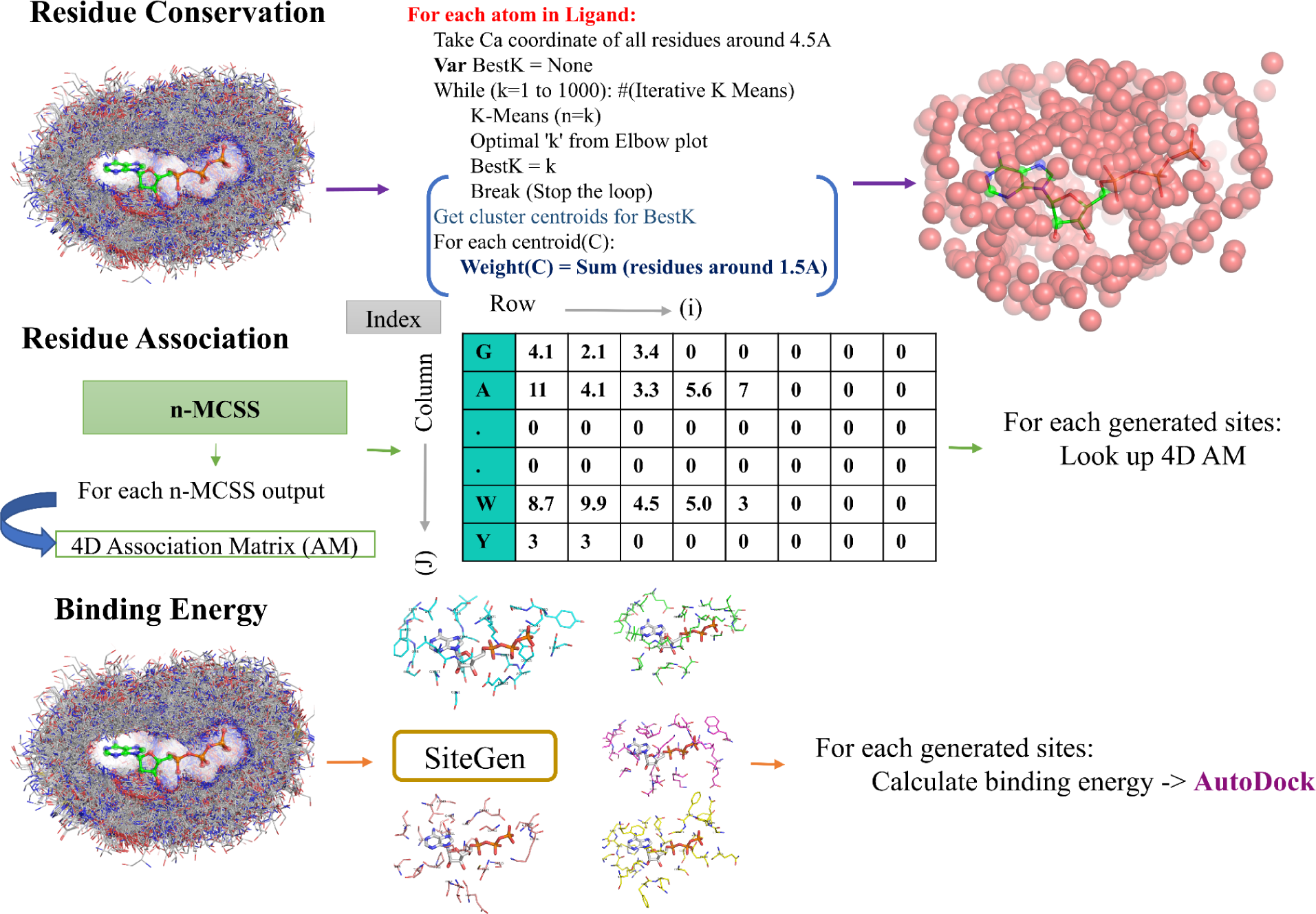
The algorithmic implementation of the triplet fitness function for estimating the quality of design sites. a) Conservation Fitness: For each atom of query ligand, we first take all residues that are present within 4.5Å. Next, we apply an iterative k-means algorithm to find the potential points for each atom. All residues present within 1.0Å from each of potential points are then considered as conserved residues. b) Residue Association Fitness: Each PDB fragment is matched after Kabsch alignment, residues in the site of the matched PDB ligand obtained and the atom-atom correspondences between two ligands extracted. A zero-value filled association matrix of shape KxLxMx10 is constructed where L equals the total atom of the query ligand and K equals the number of matched pdb fragments. For each N of M, residue count value is calculated and replaced with the corresponding cells in the AM table. c) Energy Fitness: Calculating the binding energy using AutoDock.

**Figure S4.**
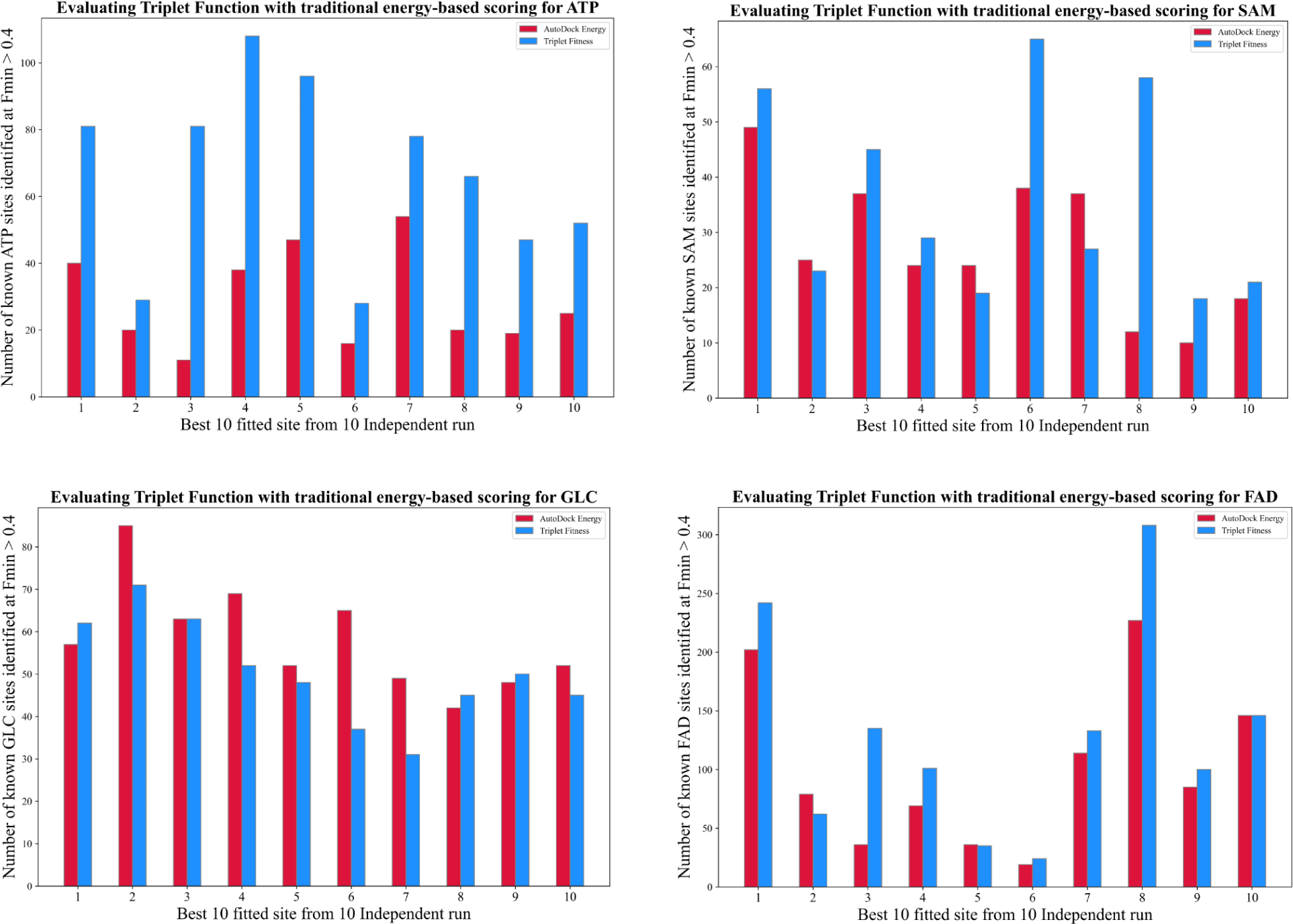
Importance of the Triplet fitness function in site design. Comparison of the accuracy of the triplet term with the AutoDock energy term alone, by comparing 10 top-scoring CRD designed sites to actual binding sites of the same ligand in PDB. For each independent iteration, the SiteGen module generates 100 sites, from which the ten best sites based on the Autodock or the Triplet functions are chosen independently. For example, in the case of an ATP ligand, top 10 sites for each fitness were compared to known ATP-bound complexes in PDB using FLAPP keeping a similarity threshold of F_min_ > 0.4. The analysis was carried out ten times. Evidently, the sites selected by our fitness method resulted in many more good matches than those selected merely by energy.

**Figure S5.**
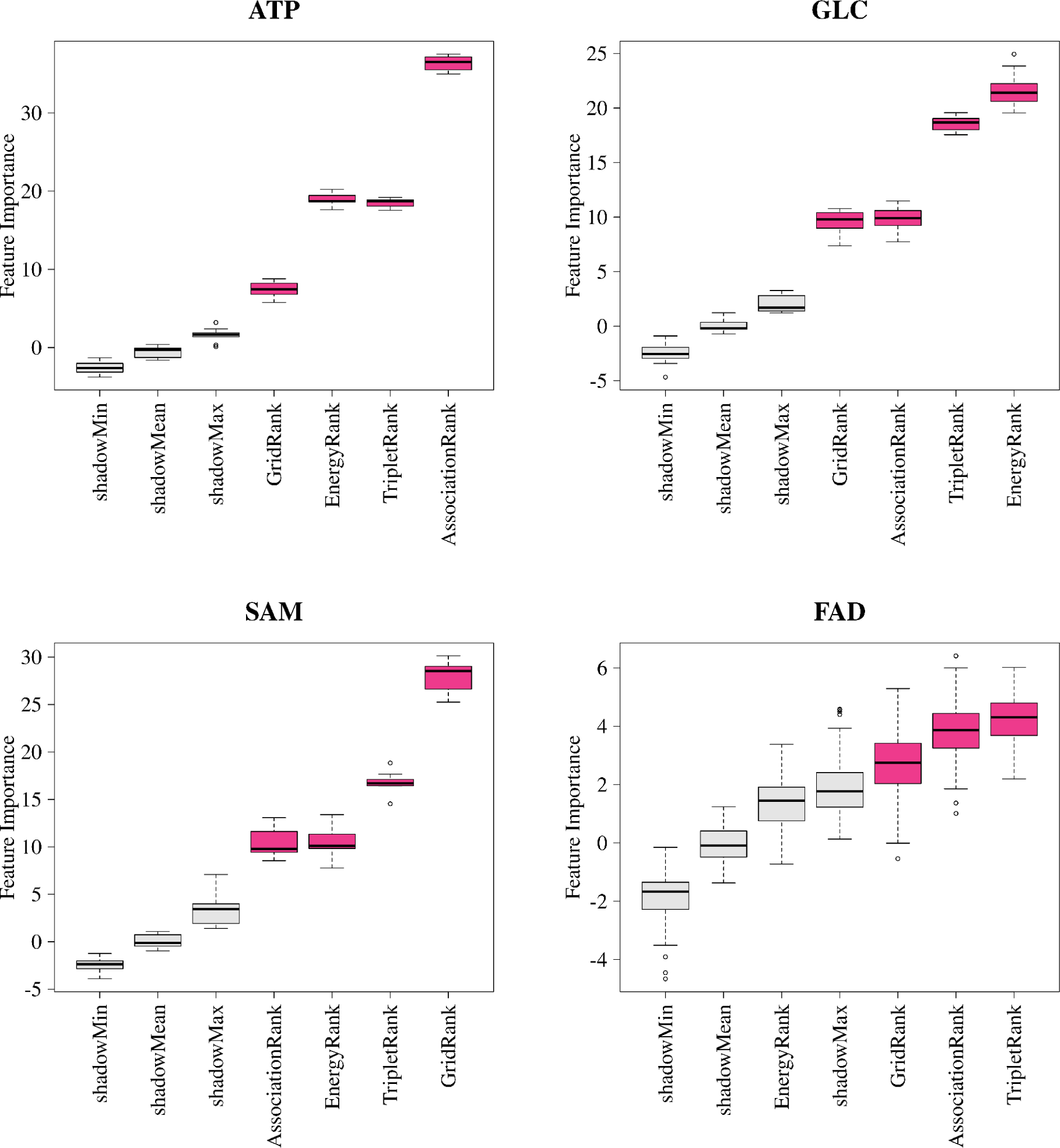
Estimation of the relevance of each of three fitness functions, Using the Boruta algorithm. SiteGen generated 100 sites for each ligand, of which the top 10 were selected based on feature importance. For each of the 10 sites, the total number of matches with known binding sites in PDB was calculated. The triplet rank is seen to be among the top 2 ranks as an important feature in selecting the most promising 10 sites from 100 initial sites across all four ligands.

**Figure S6.**
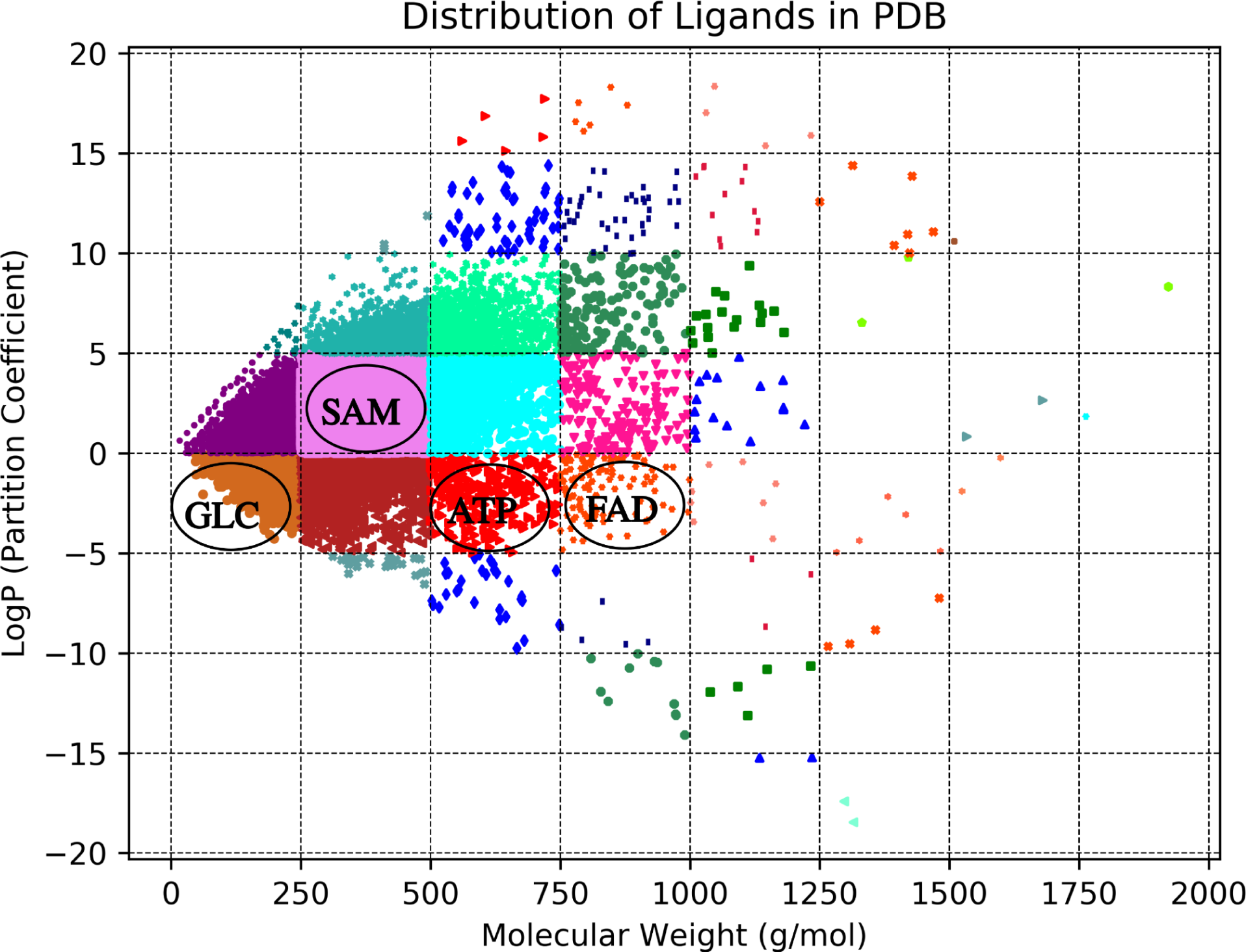
A Scatter plot illustrating the distribution of ligand molecules present in the PDB. For every ligand, two chemical descriptors were computed: 1) the molecular weight and 2) the partition coefficient. LogP is a direct correlation between ligands and solubility. The higher the LogP the more lipophilic the ligand. From the descriptors, a spaced 2D bin was created, from which four distinct ligands, one from each grid, were chosen for sensitivity testing. The ligands used for this study are ATP(Adenosine-TriPhosphate), SAM(S-Adenosylmethionine), FAD(Flavin-Adenine Dinucleotide) and GLC(Glucose).

**Figure S7.**
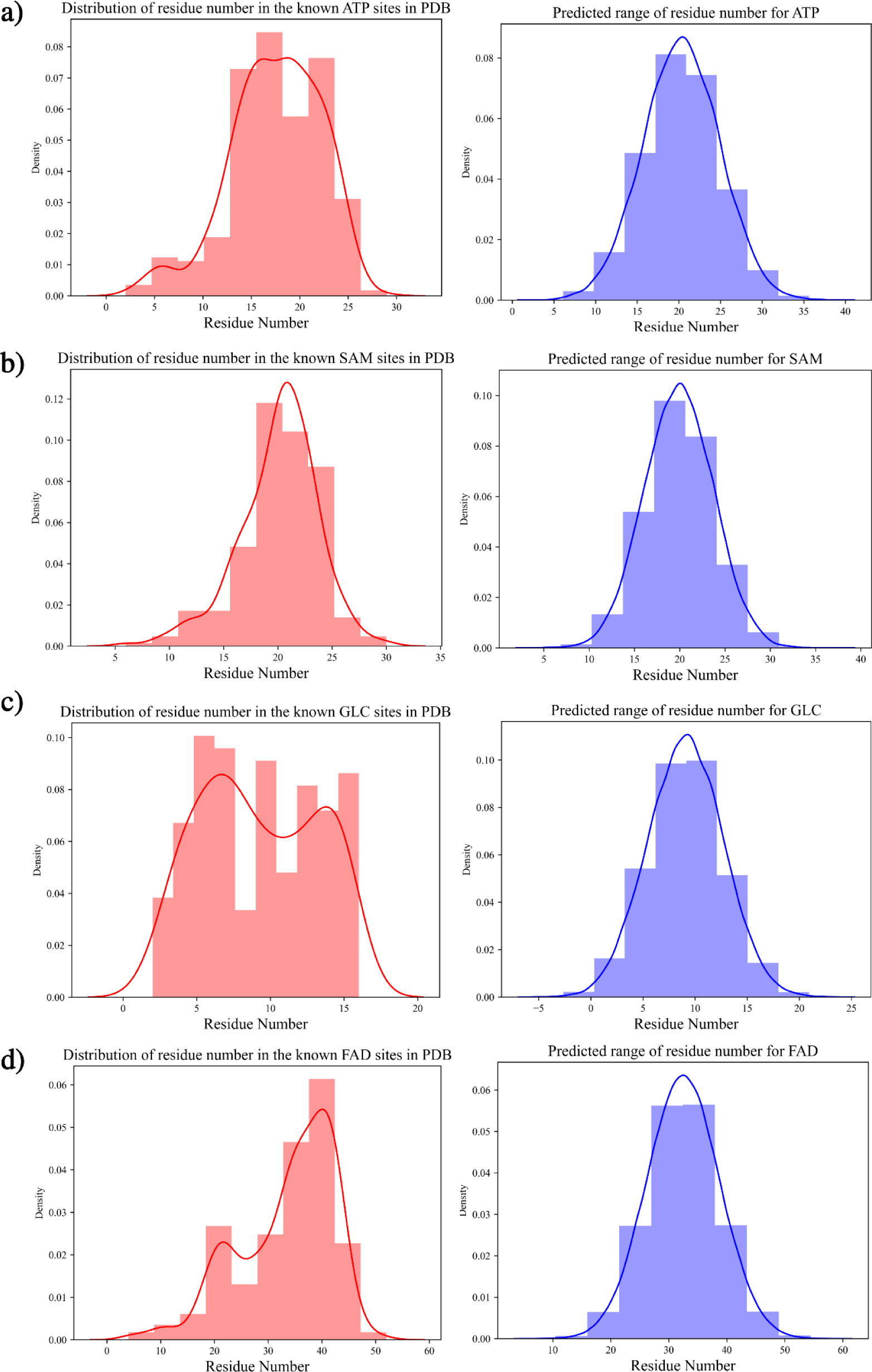
Comparing the binding site size distribution of known sites to the predicted probabilities. The RDkit software was used to extract 451 features from each subject ligand, which were then used to construct a probabilistic model based on Tensorflow. As a loss function, negative log likelihood was utilised, which was optimised using the Adam optimiser with a learning rate of 0.01. The distributions predicted for all four ligands overlap with the actual range.

**Figure S8.**
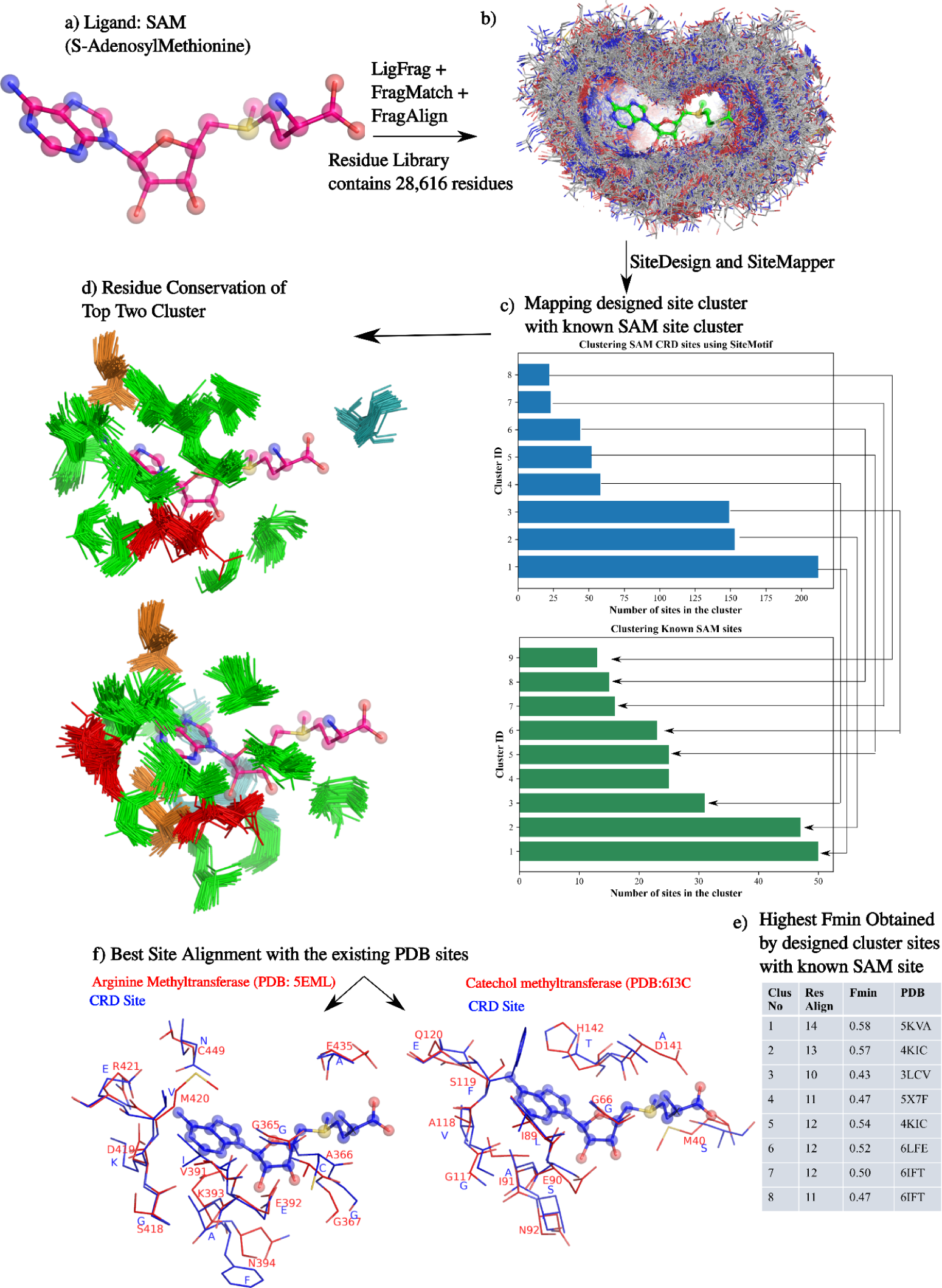
Design of binding sites for (a) SAM. b) FragMatch identifies ten fragments that, when aligned to the query ligand, result in a residue-library of 100 amino acids. c) SiteDesign and SiteMapper modules were successively implemented which grouped 750 designed sites into 8 clusters with each one of them sharing similarity with 8 known SAM types. e) Comparing sites of all designed cluster members revealed that all eight clusters have at least one member sharing good site alignment with known SAM binding site (F_min_ > 0.4). f) Finally, the high scoring pairwise alignment was identified by comparing 750 sites to 689 known SAM sites, which found two known SAM binding proteins, arginine methyltransferase (PDB:5EML) and Catechol methyltransferase (PDB:6I3C) as the top two hits.

**Figure S9.**
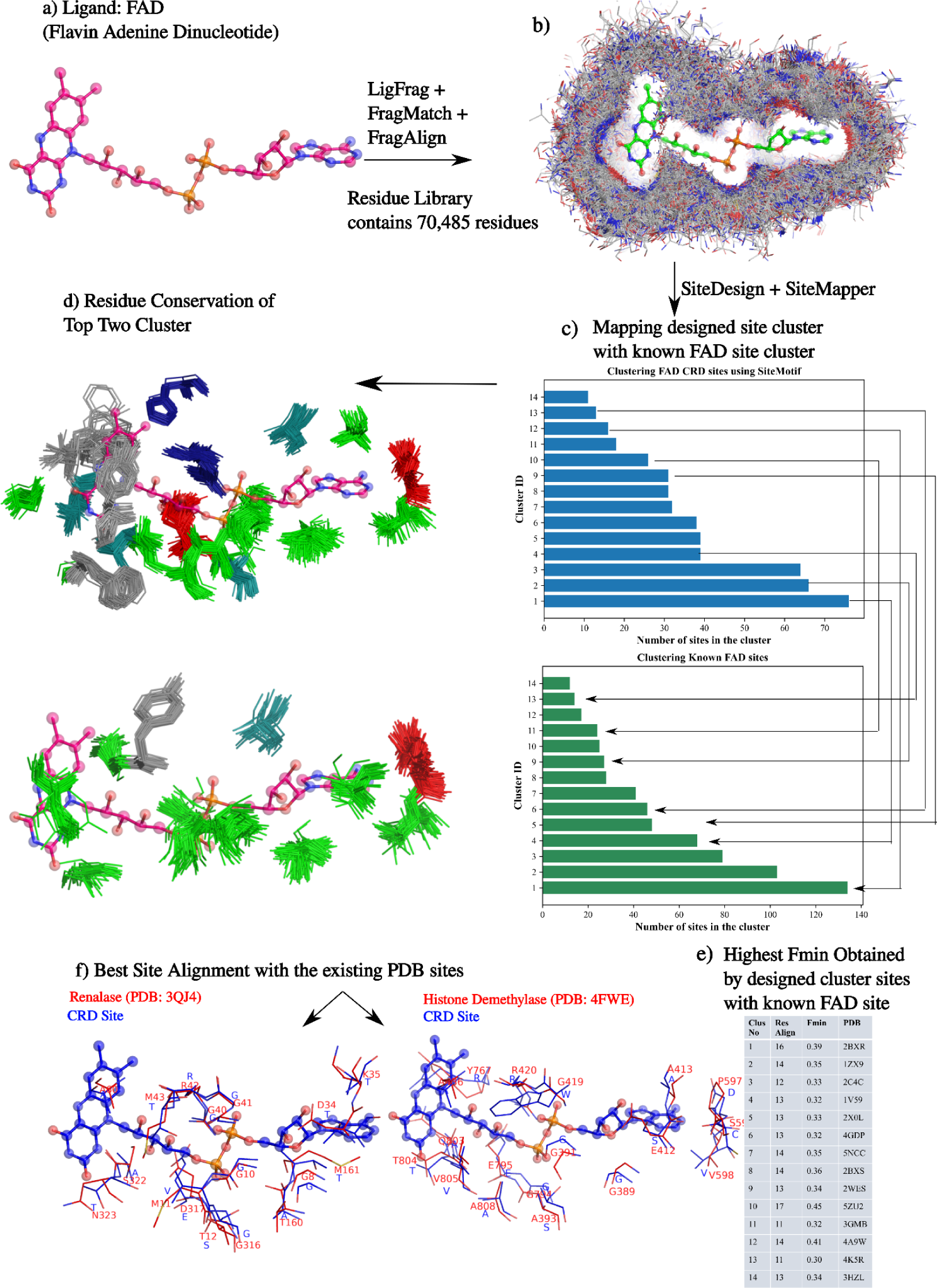
Design of binding sites for (a) FAD. b) By matching 1,66,14 fragments to the query ligand, FragAlign creates a residue-library of 70,485 amino acids. c) Following the implementation of the SiteDesign and SiteMapper modules, 750 designed sites were grouped into 14 clusters, each of which shared similarities with 7 actual FAD types. f) Among the known FAD-binding proteins, Renalase and Histone Demethylase showed the best pairwise site alignment with the designed sites.

**Figure S10.**
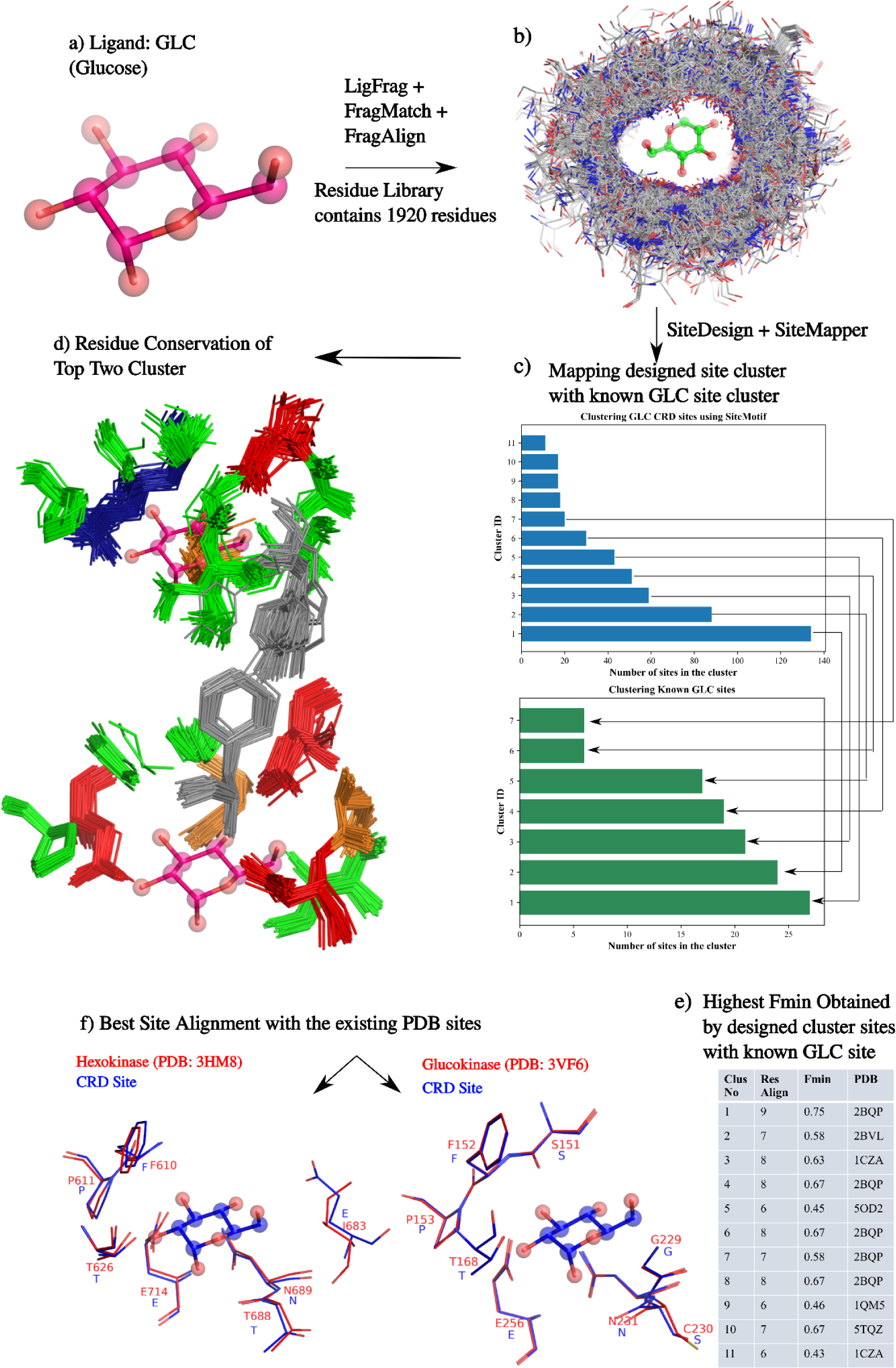
Design of binding sites for (a) GLC ligand. (b) FragAlign created a residue-library of 11,393 amino acids surrounding the glucose molecule. (c) Using the 750 designed sites, SiteClust grouped them into 11 clusters, of which 6 of them share commonality with the 7 known glucose binding site types; (d) SiteMotifs of the top two clusters, (e) best scoring hits for each of the clusters and (f) pairwise alignments with the best scoring hits hexokinase and glucokinase are shown.

**Figure S11.**
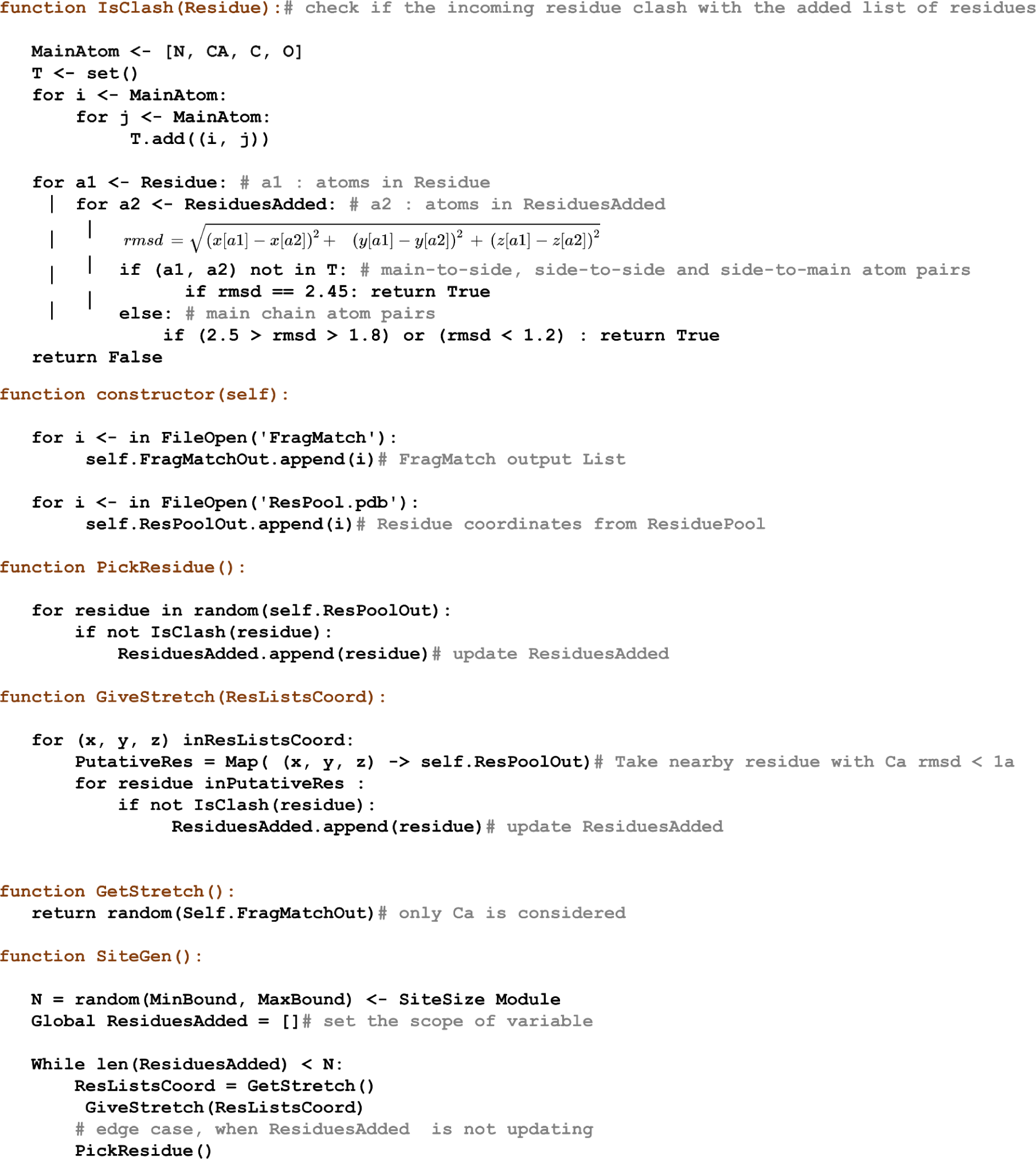
SiteGen Pseudocode.

**Figure S12.**
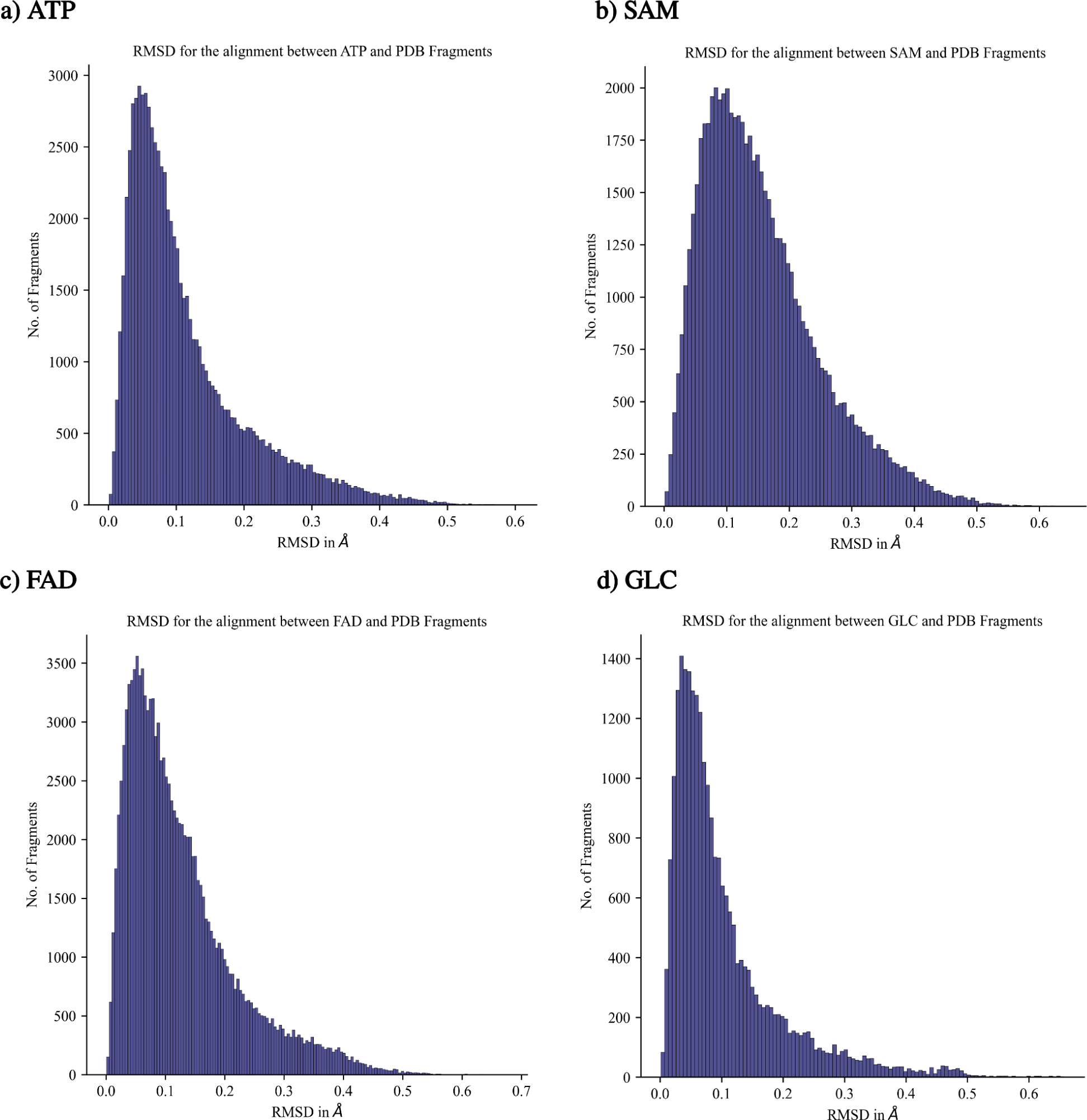
RMSD distribution obtained for fragments by matching PDB-Ligand to the query. For all four ligands, the FragMatch function identifies a number of common fragments which are then aligned using FragAlign. It was observed that the majority of aligned fragments have an RMSD < 0.2Å indicating that the deviation is negligible and FragAlign works as intended.

**Figure S13.**
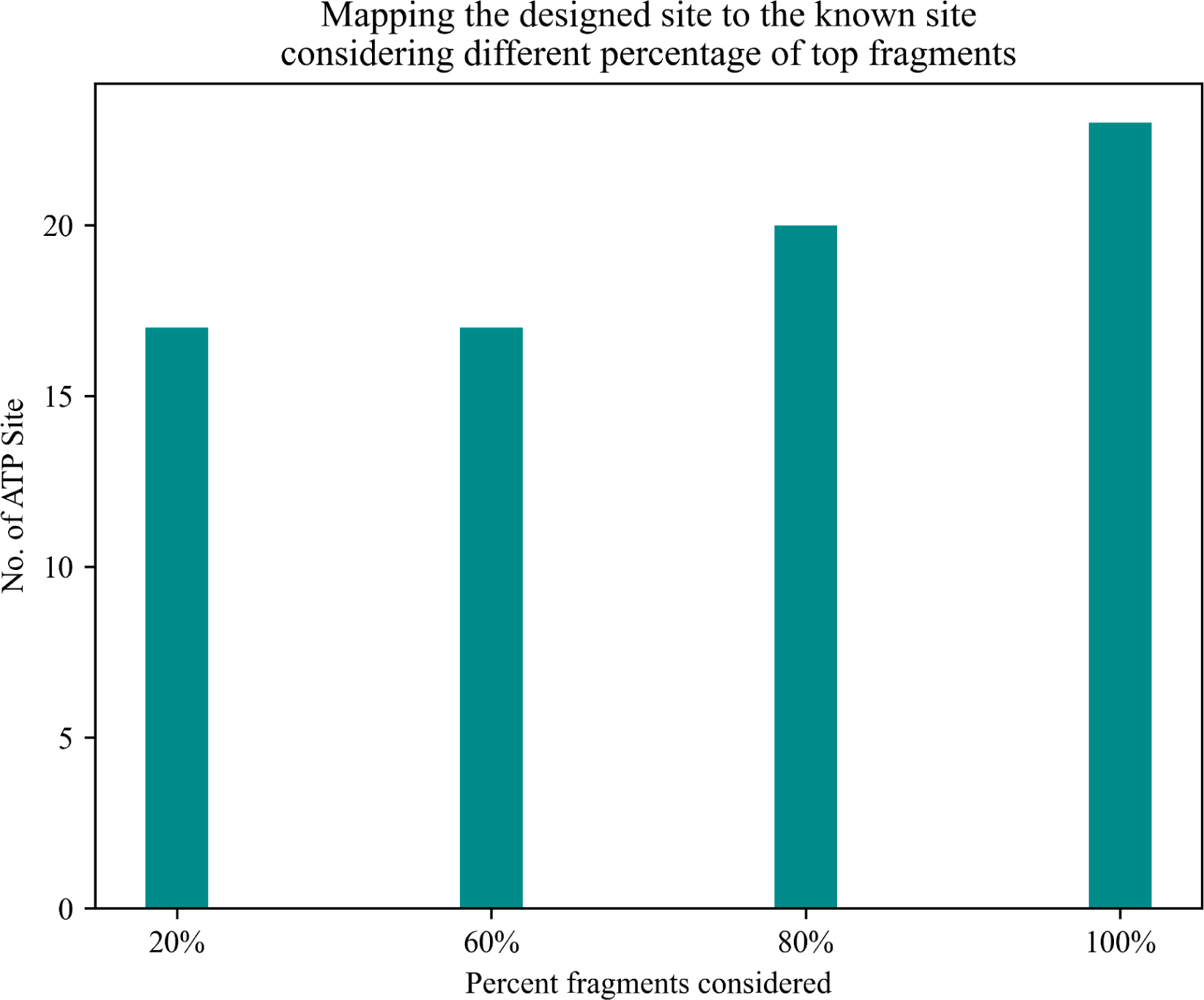
Comparing SiteDesign success rates based on only the top few proportions of fragment results. The FragMatch program outputs a list of PDB fragments aligned using FragAlign, and the top subsets are pruned using LibGen. A total of 20 sites were generated for ATP, whose similarity was compared with known ATP sites. It was necessary to perform this analysis since the RAM utilization for successive CRD functions exceeded 64 GB (memory overflow). As a result, only the top 20% of fragments were taken. This enables seamless execution of the entire CRD pipeline on a typical desktop computer. Our CRD pipeline ensured that the correct site was generated even when only 20% of the fragments were considered.

**Figure S14.**
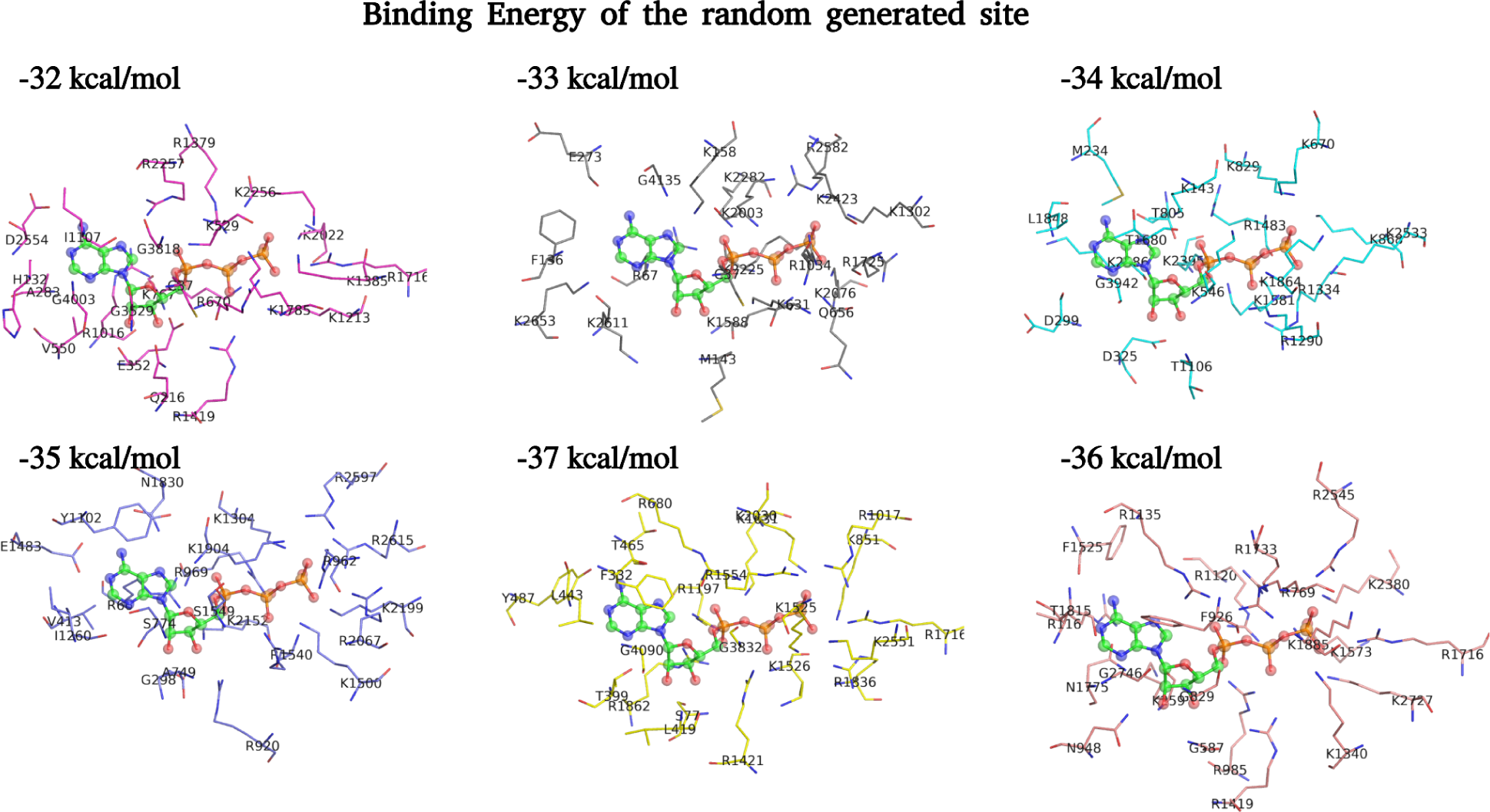
Predicting known binding site geometry using an energy-based function. Using random residue placement, we generated an initial set of six decoy sites, which were tuned by mutating portions of them. If the binding energy of the mutated site is greater than that of the decoy, then the mutated site will be used as the decoy in the subsequent iteration. This step was repeated ten times and the final site was inspected. Our observations showed that all 6 sites contained 5 to 7 lysines near phosphate and had a binding energy of -34 kcal/mol. Regardless, there is no resemblance between any of the six sites and the known ATP sites. Also, the average binding energy at the known ATP binding site is -23 kcal/mol. A clear case of overfitting was observed when selection happened based on the energy.

**Figure S15.**
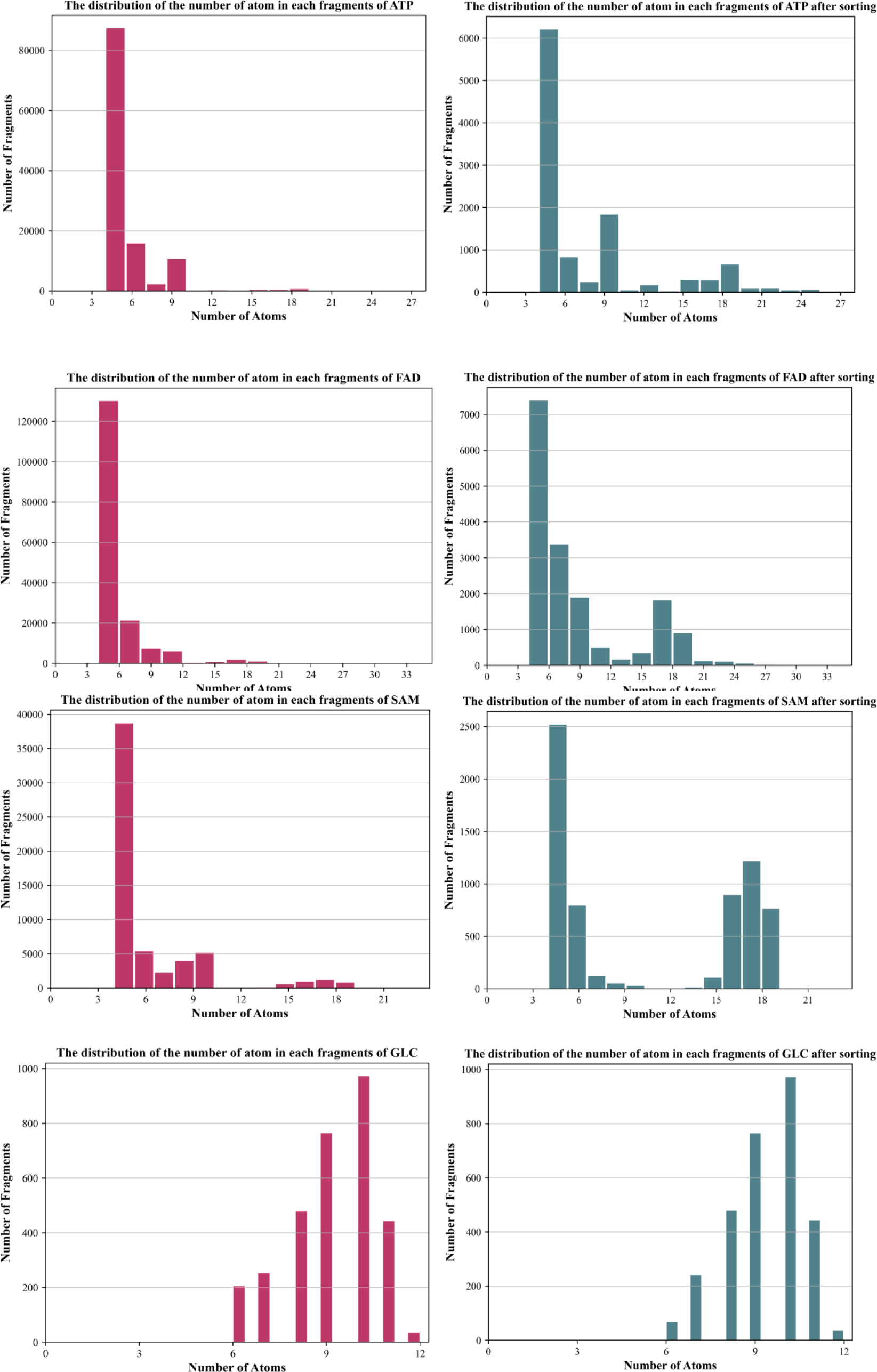
Distribution of the number of atoms present in each matched fragment for each ligand under study.

**Table S1.** Identification of potential targets for penicillin class of drugs. Sites generated for all four penicillin drugs were compared against NRSiteDB using FLAPP maintaining a high threshold of F_min_ > 0.7. In addition to identifying known targets such as penicillin binding proteins and beta-lactamase as probable binders, the site comparison exercise uncovered 37 new proteins that could potentially recognise penicillin drugs. Docking exercises were conducted to validate each protein’s theoretical affinity.

| S.No | PDB | Prot Name | AIC | MII | PNN | AXL |
| --- | --- | --- | --- | --- | --- | --- |
| **1** | **6KGV_A**<br>**(control)** | **Penicillin-binding protein** | **-6** | **-5.43** | **-7** | **-7.9** |
| **2** | **3N7W_A**<br>**(control)** | **Beta Lactamase** | **-6.7** | **-5.81** | **-7.41** | **-8.82** |
| **3** | 1F5N_A | INTERFERON INDUCED GUANYLATE BINDING PROTEIN 1 | -10.43 | -9.45 | -10.89 | -9.81 |
| **4** | 2XFU_A | Amine oxidase flavin containing B | -8.73 | -7.86 | -9.44 | -10.64 |
| **5** | 2X0L_A | LYSINE SPECIFIC HISTONE DEMETHYLASE 1 | -8.8 | -9.16 | -8.86 | -9.47 |
| **6** | 2CFY_A | THIOREDOXIN REDUCTASE 1 | -9.56 | -7.71 | -8.97 | -9.95 |
| **7** | 3CUK_A | D amino acid oxidase | -8.47 | -8.36 | -9.12 | -9.77 |
| **8** | 3T42_A | Aldose reductase | -8.19 | -7.98 | -8.9 | -10.5 |
| **9** | 1GNP_A | C H RAS P21 PROTEIN | -8.3 | -8.44 | -8.55 | -9.72 |
| **10** | 4FWE_A | Lysine specific histone demethylase 1B | -8.62 | -8.37 | -9.08 | -8.75 |
| **11** | 6MNX_A | GTPase KRas | -7.25 | -7.46 | -10.61 | -8.85 |
| **12** | 6RKF_A | D aspartate oxidase | -8.38 | -7.78 | -8.64 | -8.99 |
| **13** | 4UV9_A | LYSINE SPECIFIC HISTONE DEMETHYLASE 1A | -8.46 | -7.64 | -8.82 | -8.43 |
| **14** | 4CQM_A | CARBONYL REDUCTASE FAMILY MEMBER 4___ESTRADIOL 17 BETA DEHYDROGENASE 8 | -9.09 | -6.95 | -7.91 | -9.35 |
| **15** | 1O00_A | Aldehyde dehydrogenase | -7.75 | -7.62 | -8.9 | -8.94 |
| **16** | 5AC2_A | RETINAL DEHYDROGENASE 1 | -8.72 | -6.72 | -8.54 | -8.95 |
| **17** | 3QJ4_A | Renalase | -8.98 | -6.91 | -7.92 | -9.1 |
| **18** | 5AP4_A | DUAL SPECIFICITY PROTEIN KINASE TTK | -7.84 | -7.84 | -7.91 | -9.08 |
| **19** | 3UOK_A | Aurora kinase A | -7.94 | -8 | -8.18 | -8.53 |
| **20** | 4P4S_B | Interferon induced GTP binding protein Mx1 | -8.57 | -5.91 | -9.73 | -8.39 |
| **21** | 5GTY_A | Epidermal growth factor receptor | -7.74 | -6.6 | -9.05 | -8.95 |
| **22** | 6W0Z_A | Ketohexokinase | -8.77 | -6.3 | -7.24 | -9.96 |
| **23** | 1ZMD_A | Dihydrolipoyl dehydrogenase | -8.37 | -6.04 | -7.96 | -9.62 |
| **24** | 7L0F_A | GTPase HRas___Monobody 12VC3 | -7.35 | -6.58 | -8.58 | -9.4 |
| **25** | 1U3T_A | Alcohol dehydrogenase alpha chain | -7.47 | -6.87 | -8.07 | -9.1 |
| 26 | 3LXK_A | Tyrosine protein kinase JAK3 | -7.53 | -6.99 | -8.13 | -8.83 |
| 27 | 3L8P_A | Angiopoietin 1 receptor | -7.85 | -7.02 | -8.06 | -8.47 |
| 28 | 6BBU_A | Tyrosine protein kinase JAK1 | -7.98 | -6.42 | -8.21 | -8.71 |
| 29 | 4K81_B | GTPase HRas____Growth factor receptor bound protein 14 | -7.33 | -7.21 | -8.15 | -8.62 |
| 30 | 3Q5H_A | Renin | -8.68 | -5.8 | -7.34 | -9.24 |
| 31 | 3NAY_A | 3 phosphoinositide dependent protein kinase 1 | -7.77 | -7.18 | -7.39 | -8.6 |
| 32 | 4Y73_A | Interleukin 1 receptor associated kinase 4 | -8.28 | -5.77 | -7.29 | -8.85 |
| 33 | 6RQ4_A | Mitogen activated protein kinase 1 | -8.12 | -6.2 | -7.07 | -8.01 |
| 34 | 3FMK_A | Mitogen activated protein kinase 14 | -8.75 | -6.03 | -6.78 | -7.79 |
| 35 | 2OU7_A | Serine threonine protein kinase PLK1 | -6.67 | -7 | -7.72 | -7.88 |
| 36 | 6DBM_A | Non receptor tyrosine protein kinase TYK2 | -7.62 | -5.99 | -6.88 | -8.61 |
| 37 | 5H0B_A | Tyrosine protein kinase HCK | -8.06 | -5.78 | -7.14 | -7.62 |
| 38 | 3H10_A | Serine threonine protein kinase 6 | -7.03 | -6.64 | -7.01 | -7.73 |
| 39 | 5OOR_A | Serine threonine protein kinase Chk1 | -6.43 | -5.35 | -6.52 | -7.65 |

**Table S2.** Identification of potential targets for NSAID family drugs. Sites generated for all three drugs were compared against NRSiteDB using FLAPP maintaining a high threshold of F_min_ > 0.7. In addition to identifying known targets such as COX-1 / COX-2 as a probable binder, the site comparison exercise uncovered 55 new proteins that could potentially recognise this drug family. Docking exercises were conducted to validate each protein’s theoretical affinity.

| S.No | PDB | Prot Name | CEL | IBP | NPS |
| --- | --- | --- | --- | --- | --- |
| 1 | 3KK6_A<br>(control) | Prostaglandin G H synthase 1 | -9.96 | -6.25 | -6.97 |
| 2 | 5JW1_A<br>(control) | Prostaglandin G H synthase 2 | -9.95 | -6.12 | -6.62 |
| 3 | 6SQ2_A | Ras related protein Rab 8A | -9.82 | -9.38 | -9.69 |
| 4 | 5X9S_A | GTPase HRas | -10.13 | -8.83 | -9.33 |
| 5 | 5KJZ_A | cAMP dependent protein kinase type I alpha regulatory subunit | -10.28 | -8.49 | -9.13 |
| 6 | 6IYB_A | Ras related protein Rab 7a | -9.52 | -8.95 | -9.37 |
| 7 | 3RAP_R | PROTEIN G protein RAP2A | -10.06 | -8.53 | -9.21 |
| 8 | 5SZH_B | MICAL C terminal like protein | -9.79 | -8.71 | -9.24 |
| 9 | 1T91_A | Ras related protein Rab 7 | -9.72 | -8.79 | -9.16 |
| 10 | 4XOI_A | Transforming protein RhoA | -8.56 | -8.89 | -9.77 |
| 11 | 4FWE_A | Lysine specific histone demethylase 1B | -9.22 | -8.74 | -9.2 |
| 12 | 5UWO_A | GTP binding nuclear protein Ran | -8.1 | -9.3 | -9.64 |
| 13 | 6MNX_A | GTPase KRas | -8.75 | -8.77 | -9.31 |
| 14 | 5UWW_A | GTP binding nuclear protein Ran | -8.04 | -9.01 | -9.75 |
| 15 | 6A3B_A | GTP binding nuclear protein Ran | -7.7 | -9.12 | -9.89 |
| 16 | 1Z2C_A | Rho related GTP binding protein RhoC | -9.59 | -8.09 | -9.01 |
| 17 | 6ZIZ_A | GTPase NRas | -7.91 | -8.98 | -9.65 |
| 18 | 6SGE_A | Rho related GTP binding protein RhoB | -9.01 | -8.14 | -9.29 |
| 19 | 7BPH_A | Guanine nucleotide binding protein G s subunit alpha isoforms short | -8.51 | -8.73 | -9.12 |
| 20 | 5LWP_A | Nuclear receptor ROR gamma | -9.94 | -7.87 | -8.41 |
| 21 | 1NVU_Q | Son of sevenless protein homolog 1 | -8.09 | -8.53 | -9.18 |
| 22 | 2ODB_A | Human Cell Division Cycle 42 CDC42 | -8.89 | -8.3 | -8.59 |
| 23 | 4P4S_B | Interferon induced GTP binding protein Mx1 | -8.04 | -8.56 | -8.91 |
| 24 | 1QRA_A | TRANSFORMING PROTEIN P21 H RAS 1 | -9.22 | -7.82 | -8.18 |
| 25 | 2YBE_A | PHOSPHOGLYCERATE KINASE 1 | -7.92 | -8.08 | -9.2 |
| 26 | 3T42_A | Aldose reductase | -8 | -8.27 | -8.81 |
| 27 | 1ZSX_A | Voltage gated potassium channel beta 2 subunit | -8.24 | -8.07 | -8.63 |
| 28 | 4UJ3_A | RAS RELATED PROTEIN RAB 11A | -8.41 | -8.03 | -8.35 |
| 29 | 3V2B_A | Poly ADP ribose polymerase 15 | -8.01 | -7.84 | -8.82 |
| 30 | 2XFU_A | Amine oxidase flavin containing B | -8.93 | -7.52 | -8.07 |
| 31 | 5HIE_A | Serine threonine protein kinase B raf | -7.48 | -7.87 | -8.89 |
| 32 | 5EE5_B | ADP ribosylation factor like protein 1 | -7.94 | -7.76 | -8.5 |
| 33 | 2D7C_A | Ras related protein Rab 11A | -8.2 | -7.82 | -8.1 |
| 34 | 2FV5_A | ADAM 17 | -8.88 | -7.09 | -7.89 |
| 35 | 3LGP_A | Disintegrin and metalloproteinase domain containing protein 17 | -8.96 | -7.06 | -7.82 |
| 36 | 1ZMD_A | Dihydrolipoyl dehydrogenase | -9.17 | -7.01 | -7.59 |
| 37 | 3W8Q_A | Dual specificity mitogen activated protein kinase kinase 1 | -8.77 | -7.2 | -7.7 |
| 38 | 6H2V_A | Methyltransferase like protein 5 | -7.95 | -6.84 | -8.54 |
| 39 | 6RKF_A | D aspartate oxidase | -8.08 | -7.32 | -7.84 |
| 40 | 4CQM_A | ESTRADIOL 17 BETA DEHYDROGENASE 8 | -8.67 | -6.69 | -7.71 |
| 41 | 3M42_A | MAP kinase activated protein kinase 2 | -8.66 | -6.58 | -7.77 |
| 42 | 5IXS_A | L lactate dehydrogenase A chain | -7.7 | -7.52 | -7.72 |
| 43 | 5ZXB_A | Activated CDC42 kinase 1 | -7.86 | -7.1 | -7.79 |
| 44 | 6V66_A | Epidermal growth factor receptor | -8.39 | -7.16 | -7.17 |
| 45 | 3CUK_A | D amino acid oxidase | -8.28 | -6.71 | -7.73 |
| 46 | 5AC2_A | RETINAL DEHYDROGENASE 1 | -8.45 | -6.49 | -7.76 |
| 47 | 1NF7_A | Inosine 5 monophosphate dehydrogenase 2 | -8.01 | -6.95 | -7.72 |
| 48 | 1C5W_B | PROTEIN UROKINASE TYPE PLASMINOGEN ACTIVATOR | -8.1 | -6.91 | -7.67 |
| 49 | 5KPT_A | Pantothenate kinase 3 | -6.25 | -7.84 | -8.47 |
| 50 | 2BLE_A | GMP REDUCTASE I | -7.97 | -7.13 | -7.42 |
| 51 | 6Q21_A | C H RAS P21 PROTEIN CATALYTIC DOMAIN | -7.74 | -7.25 | -7.52 |
| 52 | 6M95_A | Mitogen activated protein kinase 14 | -8.74 | -6.51 | -7.25 |
| 53 | 3VAP_A | Aurora kinase A | -7.08 | -7.19 | -8.2 |
| 54 | 5FHZ_A | Aldehyde dehydrogenase family 1 member A3 | -8.03 | -7.15 | -7.21 |
| 55 | 5UY6_A | Calcium calmodulin dependent protein kinase kinase 2 | -8.86 | -6.38 | -7.07 |
| 56 | 2D1N_A | Collagenase 3 | -8.5 | -6.35 | -7.37 |
| 57 | 7KCD_A | Estrogen receptor | -8.83 | -6.71 | -6.62 |

## Supplementary Texts

### Supplementary-Text-1 (Fragment Sorting)

For each ligand, the LigFrag function generates fragments of different lengths. By default, we only consider fragments with more than four atoms. The distribution of various fragments identified for the target ligand is shown in Fig. S15. Fragments with fewer atoms are more abundant than those with more atoms. However, the larger fragment carries far more information necessary to construct a realistic binding site than the smaller fragment. In the case of ATP, the largest fragment obtained was ADP and AMP. We also noticed that, for ATP, the adenine ring had the greatest number of matches followed by ribose and tri-phosphate. As a result, a sorting step was incorporated for each atom of ligands to give preference for larger fragments, and to ensure even distribution of fragment count across ligand atoms. Thus to validate the accuracy of sorting, we applied SiteGen to generate 100 sites and selected 20 most relevant sites based on our triplet function. Using FLAPP, the selected 20 sites were compared with known ATP sites in PDB to determine how many of the known sites were accurately mapped by the designated site (Fmin > 0.4). Fig. S13 shows the recovery of the number of true sites by the designed one considering different percentages of fragments. In the top 20% of fragments, the designed site matches 17 existing ATP sites. This is smaller than the results for 100% fragments, which match 22 known sites. Unfortunately, taking into account all of the fragments during the design process results in a massive 68 GB memory load. This is significantly reduced when only 20% of the total fragment is used, as SiteDesign takes only 8GB of RAM.

### Supplementary-Text-2 (SiteGen Module)

The SiteGen Module aims at generating binding sites for the ligand. Results of FragMatch, SiteSize and Residue-library are used to accomplish this. FragMatch returns alignable fragments from PDB ligands that were in common to query fragments. For each obtained fragment, we also obtain information about the surrounding residues. When the length of total number of fragments is f and for each fragment, if the number of surrounding residues is r. Then, we get a total of fxr data. Every time SiteGen iterates, it picks a random residue list, say ‘rl’, from fxr and extracts Cα coordinates. The extracted Cα coordinates of ‘rl’ were then compared with the Cα coordinate of all residues in Residue-library and all residues of Residue-library with RMSD less than 1Å from ‘rl’ were stored in a separate variable called ‘Storer’. Afterwards, we replace the original residue present in the initial residue list ‘rl’ by the nearest residue present in ‘Storer’ via random sampling. This allows us to generate multiple initial seed sites with subsequent variability among themselves. The source code of SiteGen is shown in Fig. S11.

### Supplementary-Text-3 (Importance of triplet function)

To better comprehend this concept, consider the case of generating sites for ATP. It was observed during the design of the ATP sites that the atoms present in the triphosphate region (negatively charged) interact with asparagine and lysine (positively charged) while neglecting contributions from other amino acids. The potential energy of the site determined by the autodock improves as more positively charged residues are added. At the end of the process, we were able to compactly place five to seven positively charged residues around the triphosphate region. The autodock energies of known ATP binding sites were calculated as a baseline. The designed sites possess a mean binding energy of -34 kcal/mol, while binding energies of known site complexes range from -21 kcal/mol to -24 kcal/mol. Upon inspecting the known ATP sites, we found that no more than 2 positively charged residues were found near phosphates. Also other residues such as glycine were not present in the designed site since it contributed very little energetically. However, glycine plays a key role in the Walker motif, which is a signature for ATP recognition. While potential energies are informative and should be taken into account wherever possible, overfitting is clearly evident in these situations. To achieve balance, we devise novel fitness functions that act in tandem with the calculation of potential energy. It is worth noting here that while this new fitness term is not entirely accurate, it outperforms optimization solely based on potential energy.

### Supplementary-Text-4 (The design of binding sites for SAM ligand)

SAM: FragMatch module found 58,923 PDB-ligand fragments that were aligned to the query ligand from which 7,202 best fragments were used for further steps, generating a ‘residue library’ of 28,616 amino acids. The Res-Res module outputs 1732 k-means cluster centroids weighted as the number of residues with 1Å of each cluster and SiteSize identified site sizes for SAM to be in the range of 21 to 27 (Fig. S7b). SiteGen, coupled with the Fitness module, outputs 750 top-ranked designed sites for SAM recognition. The SiteMapper module identified that the designed sites belong to 8 clusters. The motifs from the top 2 clusters of the 750 designed sites are shown in Fig. S8d. The highest similarities with the known sites were observed in clusters 1 and 2 (Fig. S8e), having an F_min_ of 0.58 and 0.57. To check if designed sites resemble any of the known sites, we establish an unbiased site comparison exercise to compare all 750 sites against 55,301 and obtain 100 proteins at the stringent cutoff of F_min_ > 0.7. The proteins housing these known sites are identified as predicted receptors for SAM, the top-ranked ones being arginine methyltransferase and catechol methyltransferase (Fig. S8f). A docking run is performed to estimate the theoretical feasibility of that protein binding the ligand (Sheet-2 of ReceptorDocked.xlsx). Here too, as can be expected, a large number of receptors are predicted, many of which are well characterized SAM binding proteins.

### Supplementary-Text-5 (The design of binding sites for FAD ligand)

FAD: FAD is a very big ligand with 53 atoms. With FAD as the query, the FragMatch module found 1,68,289 PDB-ligand fragments that were aligned to the query ligand from which 31,169 best fragment were used for further steps, generating a ‘residue library’ of 70,485 amino acids. The Res-Res module outputs 4532 k-means cluster centroids weighted as the number of residues with 1Å of each cluster and SiteSize identified site sizes for FAD to be in the range of 32 to 39 residues (Fig. S7c). SiteGen, coupled with the Fitness module, outputs 750 top-ranked designed sites for FAD recognition. The SiteMapper module identified that the designed sites belong to 14 clusters. We did observe similarities of the designed sites matched with known FAD binding sites (SFig.9f). The proteins housing these known sites are identified as predicted receptors (Sheet-3 of ReceptorDocked.xlsx). Here too, as can be expected, a large number of receptors are predicted, many of which are well characterized FAD binding proteins.

### Supplementary-Text-6 (The design of binding sites for GLC ligand)

Glucose: Glucose (GLC) is a small highly polar ligand. For GLC as the query ligand, the FragMatch module found 3128 PDB-ligand fragments that were aligned to the query ligand from which all fragments were used for further steps, generating a ‘residue library’ of 10,307 amino acids. The Res-Res module outputs 1332 k-means cluster centroids weighted as the number of residues with 1Å of each cluster and SiteSize identified site sizes for GLC to be in the range of 10 to 15 residues (Fig. S7d). SiteGen, coupled with the Fitness module, outputs 750 top-ranked designed sites for GLC recognition. The SiteMapper module identified that the designed sites belong to 11 clusters. The proteins housing these known sites are identified as predicted receptors for GLC, the top-ranked ones being methyltransferase Hexokinase and Glucokinase. A docking run is performed to estimate the theoretical feasibility of that protein binding the ligand (Sheet-4 of ReceptorDocked.xlsx).

## References

[1] R. Bhagavat, S. Sankar, N. Srinivasan, N. Chandra, An Augmented Pocketome: Detection and Analysis of Small-Molecule Binding Pockets in Proteins of Known 3D Structure, Struct. Lond. Engl. 1993. 26 (2018) 499–512.e2. https://doi.org/10.1016/j.str.2018.02.001.

[2] S.K. Burley, H.M. Berman, G.J. Kleywegt, J.L. Markley, H. Nakamura, S. Velankar, Protein Data Bank (PDB): The Single Global Macromolecular Structure Archive, Methods Mol. Biol. Clifton NJ. 1607 (2017) 627–641. https://doi.org/10.1007/978-1-4939-7000-1_26.

[3] L. Lo Conte, B. Ailey, T.J. Hubbard, S.E. Brenner, A.G. Murzin, C. Chothia, SCOP: a structural classification of proteins database, Nucleic Acids Res. 28 (2000) 257–259. https://doi.org/10.1093/nar/28.1.257.

[4] A.C. Anderson, The Process of Structure-Based Drug Design, Chem. Biol. 10 (2003) 787–797. https://doi.org/10.1016/j.chembiol.2003.09.002.

[5] M. Batool, B. Ahmad, S. Choi, A Structure-Based Drug Discovery Paradigm, Int. J. Mol. Sci. 20 (2019) 2783. https://doi.org/10.3390/ijms20112783.

[6] C. Acharya, A. Coop, J.E. Polli, A.D. Mackerell, Recent advances in ligand-based drug design: relevance and utility of the conformationally sampled pharmacophore approach, Curr. Comput. Aided Drug Des. 7 (2011) 10–22. https://doi.org/10.2174/157340911793743547.

[7] M.J. O’Meara, S. Ballouz, B.K. Shoichet, J. Gillis, Ligand Similarity Complements Sequence, Physical Interaction, and Co-Expression for Gene Function Prediction, PloS One. 11 (2016) e0160098. https://doi.org/10.1371/journal.pone.0160098.

[8] R. Pearce, X. Huang, G.S. Omenn, Y. Zhang, De novo protein fold design through sequence-independent fragment assembly simulations, Proc. Natl. Acad. Sci. 120 (2023) e2208275120. https://doi.org/10.1073/pnas.2208275120.

[9] X. Pan, T. Kortemme, Recent advances in de novo protein design: Principles, methods, and applications, J. Biol. Chem. 296 (2021) 100558. https://doi.org/10.1016/j.jbc.2021.100558.

[10] J.R. Miller, S. Koren, G. Sutton, Assembly algorithms for next-generation sequencing data, Genomics. 95 (2010) 315–327. https://doi.org/10.1016/j.ygeno.2010.03.001.

[11] N. Nagarajan, M. Pop, Sequence assembly demystified, Nat. Rev. Genet. 14 (2013) 157–167. https://doi.org/10.1038/nrg3367.

[12] V.D. Mouchlis, A. Afantitis, A. Serra, M. Fratello, A.G. Papadiamantis, V. Aidinis, I. Lynch, D. Greco, G. Melagraki, Advances in de Novo Drug Design: From Conventional to Machine Learning Methods, Int. J. Mol. Sci. 22 (2021) 1676. https://doi.org/10.3390/ijms22041676.

[13] J. Adolf-Bryfogle, O. Kalyuzhniy, M. Kubitz, B.D. Weitzner, X. Hu, Y. Adachi, W.R. Schief, R.L. Dunbrack, RosettaAntibodyDesign (RAbD): A general framework for computational antibody design, PLOS Comput. Biol. 14 (2018) e1006112. https://doi.org/10.1371/journal.pcbi.1006112.

[14] C. Malisi, M. Schumann, N.C. Toussaint, J. Kageyama, O. Kohlbacher, B. Höcker, Binding pocket optimization by computational protein design, PloS One. 7 (2012) e52505. https://doi.org/10.1371/journal.pone.0052505.

[15] A.C. Stiel, M. Nellen, B. Höcker, PocketOptimizer and the Design of Ligand Binding Sites, Methods Mol. Biol. Clifton NJ. 1414 (2016) 63–75. https://doi.org/10.1007/978-1-4939-3569-7_5.

[16] J.E. Lucas, T. Kortemme, New computational protein design methods for de novo small molecule binding sites, PLOS Comput. Biol. 16 (2020) e1008178. https://doi.org/10.1371/journal.pcbi.1008178.

[17] E.S. Salmina, N. Haider, I.V. Tetko, Extended Functional Groups (EFG): An Efficient Set for Chemical Characterization and Structure-Activity Relationship Studies of Chemical Compounds, Mol. Basel Switz. 21 (2015) E1. https://doi.org/10.3390/molecules21010001.

[18] S. Sankar, N. Chandran Sakthivel, N. Chandra, Fast Local Alignment of Protein Pockets (FLAPP): A System-Compiled Program for Large-Scale Binding Site Alignment, J. Chem. Inf. Model. 62 (2022) 4810–4819. https://doi.org/10.1021/acs.jcim.2c00967.

[19] W. Kabsch, A solution for the best rotation to relate two sets of vectors, Acta Crystallogr. Sect. A. 32 (1976) 922–923. https://doi.org/10.1107/S0567739476001873.

[20] J. Dong, Z.-J. Yao, L. Zhang, F. Luo, Q. Lin, A.-P. Lu, A.F. Chen, D.-S. Cao, PyBioMed: a python library for various molecular representations of chemicals, proteins and DNAs and their interactions, J. Cheminformatics. 10 (2018) 16. https://doi.org/10.1186/s13321-018-0270-2.

[21] G.M. Morris, D.S. Goodsell, R. Huey, A.J. Olson, Distributed automated docking of flexible ligands to proteins: Parallel applications of AutoDock 2.4, J. Comput. Aided Mol. Des. 10 (1996) 293–304. https://doi.org/10.1007/BF00124499.

[22] M.B. Kursa, W.R. Rudnicki, Feature Selection with the Boruta Package, J. Stat. Softw. 36 (2010). https://doi.org/10.18637/jss.v036.i11.

[23] S. Sankar, N. Chandra, SiteMotif: A graph-based algorithm for deriving structural motifs in Protein Ligand binding sites, PLoS Comput. Biol. 18 (2022) e1009901. https://doi.org/10.1371/journal.pcbi.1009901.

[24] O. Dym, D. Eisenberg, Sequence-structure analysis of FAD-containing proteins, Protein Sci. Publ. Protein Soc. 10 (2001) 1712–1728. https://doi.org/10.1110/ps.12801.

[25] A. Narunsky, A. Kessel, R. Solan, V. Alva, R. Kolodny, N. Ben-Tal, On the evolution of protein-adenine binding, Proc. Natl. Acad. Sci. U. S. A. 117 (2020) 4701–4709. https://doi.org/10.1073/pnas.1911349117.

[26] J. Jumper, R. Evans, A. Pritzel, T. Green, M. Figurnov, O. Ronneberger, K. Tunyasuvunakool, R. Bates, A. Žídek, A. Potapenko, A. Bridgland, C. Meyer, S.A.A. Kohl, A.J. Ballard, A. Cowie, B. Romera-Paredes, S. Nikolov, R. Jain, J. Adler, T. Back, S. Petersen, D. Reiman, E. Clancy, M. Zielinski, M. Steinegger, M. Pacholska, T. Berghammer, S. Bodenstein, D. Silver, O. Vinyals, A.W. Senior, K. Kavukcuoglu, P. Kohli, D. Hassabis, Highly accurate protein structure prediction with AlphaFold, Nature. 596 (2021) 583–589. https://doi.org/10.1038/s41586-021-03819-2.

[27] M. Bartolowits, V.J. Davisson, Considerations of Protein Subpockets in Fragment-Based Drug Design, Chem. Biol. Drug Des. 87 (2016) 5–20. https://doi.org/10.1111/cbdd.12631.

[28] S.K. Lam, A. Pitrou, S. Seibert, Numba: a LLVM-based Python JIT compiler, in: Proc. Second Workshop LLVM Compil. Infrastruct. HPC, ACM, Austin Texas, 2015: pp. 1–6. https://doi.org/10.1145/2833157.2833162.

[29] V.D. Blondel, J.-L. Guillaume, R. Lambiotte, E. Lefebvre, Fast unfolding of communities in large networks, J. Stat. Mech. Theory Exp. 2008 (2008) P10008. https://doi.org/10.1088/1742-5468/2008/10/P10008.

